# Activation of KCC2 during development alleviates cognitive, behavioral, and neural excitability in adult CDKL5-deficient mice

**DOI:** 10.1101/2025.02.26.640365

**Authors:** Muhammad Nauman Arshad, Christopher Bope, Jacob S Dengler, Shu Fun Josephine Ng, Joshua Smalley, Toshiya Nishi, Zhong Zhong, Stephen J Moss, Paul A Davies

## Abstract

Cyclin-dependent kinase-like 5 (CDKL5) deficiency disorder (CDD) is a developmental and epileptic encephalopathy (DEE) characterized by severe drug-resistant epileptic disorders beginning in early childhood, along with cognitive and social impairments in later childhood and adulthood. Existing pharmacological therapies for CDD primarily focus on anti-seizure medications, which often have associated sedative side effects. In addition, there are currently no effective treatments for cognitive or behavioral impairments associated with this disorder. Postnatal development expression of CDKL5 has a similar timeline as the developmental activity of the potassium chloride co-transporter (KCC2), the maturation of which is a prerequisite for the developmental switch to fast synaptic hyperpolarizing inhibition mediated by g-aminobutyric acid type A receptors (GABA_A_R). This developmental GABA switch is determined by changes in the phosphorylation of multiple residues in KCC2. During this initial postnatal period, dramatic changes occur as major neuronal circuits are formed, laying down the initial pathways important for memory consolidation and behavioral processing. Currently, a knowledge gap exists in understanding KCC2 dysfunction in CDD.

In adult *Cdkl5* KO mice we found aberrant KCC2 phosphorylation and expression, such that KCC2 phosphorylation profile appeared immature. We examined the developmental changes in KCC2 and observed significant alterations in the phosphorylation of key residues and decreased KCC2 expression from p14 to p21. Because KCC2 loss-of-function has been strongly correlated with excessive neuronal excitation, cognitive and behavioral impairments, we examined seizure susceptibility, spatial memory, and social interaction in adult *Cdkl5* KO mice following once daily administration of the KCC2 activator (OV350), or vehicle, to infant *Cdkl5* KO mice. We found that adult *Cdkl5* KO mice are more susceptible to kainate-induced seizures, show poor sociability and deficits in spatial learning and memory compared to WT mice. Twelve days of OV350 treatment as infants (p10 to p21) prevented the development of benzodiazepine-resistant seizures and alleviated cognitive and behavioral deficits in adult *Cdkl5* KO mice. In contrast, 12 days of OV350 treatment in adult *Cdkl5* KO mice had limited ability to alleviate cognitive and behavioral deficits. In summary, this study demonstrates that enhancing KCC2 function may be a potential therapeutic target for CDD and other DEEs. However, early intervention during critical developmental windows is crucial for optimal outcomes.

## Introduction

Cyclin-dependent kinase-like 5 (Cdkl5) deficiency disorder (CDD) is a severe type of neurodevelopmental and epileptic encephalopathy (DEE) that affects 1 in 40,000 to 75,000 live births, and is one of the most common genetic forms of infantile epilepsy (Symonds and McTague, 2020 Symonds et al., 2019). CDD is characterized by early-onset treatment-resistant seizures beginning in early childhood, accompanied by severe neurodevelopment impairments and subsequent cognitive and social deficits throughout life (D’Mello, 2023; Leonard et al., 2022). Treatment-resistant seizures significantly raise the risk of injury or death (Laxer et al., 2014), highlighting the importance of understanding the underlying mechanisms and developing effective treatments. Current pharmacological therapies for CDD primarily focus on anti-seizure medications (ASM), such as the GABAergic enhancing neuroactive steroid, ganaxolone (Knight et al., 2022). However, ASMs (especially when given as polytherapy) are associated with side effects such as somnolence, which limit the quality of life (Olsen et al., 2023; Wong et al., 2024). Unfortunately, there is no effective treatment for cognitive or behavioral impairments associated with this disorder.

Most of the neuropathological changes and behavioral deficits that occur in human CDD patients can be recapitulated in constitutive *Cdkl5* knockout (KO) mouse models. Like human patients, loss of functional CDKL5 results in increased anxiety, depression, and fear-related behavior, along with an impairment in the acquisition and retention of spatial memory (Okuda et al., 2018). These *Cdkl5* KO mice also show abnormal adult neurogenesis, reduced dendritic arborization, and disruption in the organization of excitatory and inhibitory synapses (Amendola et al., 2014; Pizzo et al., 2016). Recently, a study showed a reduction in the protein levels of phosphorylated K^+^/Cl^-^ cotransporter 2 (KCC2) in the cortex of neonatal *Cdkl5* KO pups, which leads to the development of spontaneous recurrent seizures (Liao et al., 2023).

CDKL5 is a serine/threonine protein kinase and is known to be essential for normal brain development. Postnatal developmental expression of CDKL5 has a similar timeline for the developmental activity of the potassium chloride co-transporter (KCC2). During this initial postnatal period dramatic changes occur as major neuronal circuits are formed that lay down the initial pathways important for memory consolidation, sensory, and behavioral processing (Virtanen et al., 2021). KCC2 is the principal Cl^-^-extrusion mechanism employed by developing and mature neurons in the CNS (Moore et al., 2017). Its activity is a prerequisite for the efficacy of fast synaptic inhibition mediated by g-aminobutyric acid type A receptors (GABA_A_R), which are Cl^-^ permeable ligand-gated ion channels. The postnatal development of canonical hyperpolarizing GABA_A_R currents reflects the progressive decrease of intraneuronal Cl^-^ levels, caused by the upregulation of KCC2 expression and subsequent activity (Moore et al., 2019). The developmental appearance of hyperpolarizing GABA_A_R currents is determined by the phosphorylation status of KCC2, a process that facilitates its membrane trafficking and activity (Lee et al., 2007; Moore et al., 2019). Deficits in KCC2 expression levels and activity have been detailed in patient and animal models of epilepsy (Sivakumaran et al., 2015; Kelley et al., 2016). Furthermore, we, and others, have demonstrated that KCC2 loss of function is strongly correlated with cognitive impairment (in Fragile X syndrome and Rett syndrome) and the development of pharmaco-resistant seizures that are insensitive to GABA_A_R positive allosteric modulators such as benzodiazepines (Deshpande et al., 2007; Deeb et al., 2012; Duarte et al., 2013; He et al., 2014; Tang et al., 2019).

The current understanding of whether KCC2 expression, phosphorylation, and activity change during development in *Cdkl5* KO pups remains unclear. Furthermore, it is crucial to investigate whether enhancing KCC2 activity during postnatal development, when neural circuits are maturing and the transition from excitation to inhibition occurs in immature neurons, could potentially normalize behavioral and cognitive deficits in Cdkl5 KO mice. To address this issue, we have developed a novel small molecule activator, OV350, which potentiates KCC2 activity and effectively terminates pharmaco-resistant seizures in wild-type (WT) mice (Jarvis et al., 2023). Here, we demonstrated the effects of ablating CDKL5 on the phosphorylation of KCC2 and its impact on the development of pharmaco-resistant seizures and cognitive and behavioral deficits. We observed that the pharmacological activation of KCC2 during postnatal development is an important time period for an intervention to improve adult sociability and cognition. We also showed that KCC2 activity during postnatal development reduces adult baseline EEG power and restores the ability of diazepam to terminate intractable SE in *Cdkl5* KO mice.

## Methods

### Animals

The TUFTS University Institutional Animal Care and Use Committee approved all animal use. The animals were housed in temperature-controlled rooms on a 12-hour day/night cycle. We purchased the *Cdkl5* KO mice from The Jackson Laboratory (Strain #:021967) and used a mix of male and female *Cdkl5* KO mice for our experiments. See Table 1 for the key resources.

**Table 1.** Key Resources.

| Name | Company | Catalog Number |
| --- | --- | --- |
| **Animals** |  |  |
| C57BL/6J | The Jackson Laboratory | 000664 |
| 6.129(FVB)-<br>Cdkl5tm1.1Joez/J | The Jackson Laboratory | 021967 |
| **Equipment, Chemicals and Reagents** |  |  |
| OV350 (KCC2 activator) | Ovid Therapeutics |  |
| Kainic Acid | Caymon | 58002-62-3 |
| Buprenorphine SR | Wedgewood Connect | 29926-05717 |
| Diazepam | Torics | 2805/50 |
| Captisol (SBE-b-CD) | LigandPharmaceuticals | RC-0C7-020 |
| Dimethyl sulfoxide | Sigma | D8418 |
| Mouse KCC2 | NeuroMab | N1/12 |
| Mouse Actin | Sigma | A1978 |
| Rabbit beta-Tubulin | Cell Signaling | 2128 |
| Rabbit KCC2 pS940 | Phosphosolutions | p1551-940 |
| Rabbit KCC2 pT1007 | Phosphosolutions | p1551-1007 |
| Rabbit KCC2 PT906 | Phosphosolutions | p1551-906 |
| Mouse Cdkl5, clone 8F3.1 | Sigma | MABS1132 |
| Protease inhibitor | Roche | 11836170001 |
| Phosphatase inhibitor | Roche | 4906837001 |
| Protein G Dynabeads | Thermo Fisher | 10004D |
| Bradford assay | BioRad | 5000001 |
| Bovine Serum Albumin | Sigma | A7030 |
| Stereotaxic apparatus | Kopf |  |
| Isothesia (Isoflurane) | Henry Schein | SKU 029405 |
| Charcoal filter, VetEquip | Harvard Apparatus |  |
| Paraformaldehyde | Electron Microscopy Sciences | 15174-S |
| EEG/EMG headmounts | Pinnacle Technology | 8201 |
| **Software** |  |  |
| ImageJ | Schneider et al. | https://imagej.nih.gov/ij/ |
| GraphPad Prism | Prism |  |
| LabChart 8.1 | ADInstruments | https://www.adinstruments.com |
| Ethovision | Noldus | https://www.noldus.com/ethovision |

### Drug preparation

OV350 was formulated with 6.25% DMSO and 93.75% (v/v) of 50% (w/v) Captisol, and the Vehicle is 6.25% DMSO and 93.75% (v/v) of 50% (w/v) Captisol. Mice were injected with 50 mg/kg OV350, a dose which has previously been shown to reach a brain concentration of 676 nM within 4 hours, and is maintained for 8 hours (Jarvis et al., 2023).

### Immunoblotting

Sodium dodecyl sulfate-polyacrylamide gel electrophoresis (SDS-PAGE) was carried out as previously described (Smalley et al., 2020; Choi et al., 2022). Briefly, proteins were isolated in RIPA lysis buffer (50 mM Tris, 150 mM NaCl, 0.1% SDS, 0.5% sodium deoxycholate, and 1% Triton X-100, pH 7.4) supplemented with mini cOmplete protease inhibitor and PhosSTOP phosphatase inhibitor tablets. Protein concentration was measured using a Bradford assay (Bio-Rad,). Samples were diluted in 2x sample buffer, and 30 μg of protein was loaded onto a 7% polyacrylamide gel, depending on the molecular mass of the target protein. After separation by SDS-PAGE, proteins were transferred onto a nitrocellulose membrane. Membranes were blocked in 5% skimmed milk or Bovine Serum Albumin in tris-buffered saline 0.1% Tween-20 (TBS-T) for 1 h, washed with TBS-T, and then probed with primary antibodies diluted in TBS-T (dilution and incubation time dependent on the antibody). The membranes were washed and incubated for 1 h at room temperature with HRP-conjugated secondary antibodies (1:5000 – Jackson ImmunoResearch Laboratories). Protein bands were visualized with Pierce ECL (Thermo Fisher Scientific) and imaged using a ChemiDoc MP (Bio-Rad). Band intensity was compared to α-tubulin as a loading control.

### Plasma membrane isolation

Plasma membranes were isolated using a method previously described (Smalley et al., 2020; Choi et al., 2022). Fresh cortex/hippocampus tissues from 8–12 weeks old male and female mice were dissected in ice-cold 1× phosphate-buffered saline (PBS) and collected in a starting buffer (225 mM mannitol, 75 mM sucrose, and 30 mM Tris-HCl, pH 7.4) as described previously (Smalley et al., 2023). Forebrain tissues from 7 animals were pooled and homogenized in isolation buffer [225 mM mannitol, 75 mM sucrose, 0.5% BSA (w/v), 0.5 mM EGTA, and 30 mM Tris-HCl, pH 7.4] supplemented with a protease inhibitor (cOmplete mini, EDTA-free Protease Inhibitor Cocktail, Roche) and a phosphatase inhibitor (PhosSTOP, Roche) on ice using 14 strokes of a Dounce homogenizer. The homogenates were subjected to serial centrifugation to isolate purified PM and ER fractions. The PM and ER fractions were solubilized in Triton lysis buffer [150 mM NaCl, 10 mM Tris, 0.5% Triton X-100 (v/v), pH 7.5] supplemented with protease and phosphatase inhibitors.

### Immunoprecipitation

Protein G Dynabeads (Thermo Fisher) were washed three times with 1× PBS with 0.05% Tween-20 (v/v; 0.05% PBS-T). The beads were resuspended in 0.05% PBS-T and incubated overnight at 4°C with antibodies for the target protein or non-immune mouse IgG (Jackson ImmunoResearch 015-000-003). The beads were washed twice with 0.2 M triethanolamine (TEA; pH 8.2) and incubated for 30 min with 40 mM dimethyl pimelimidate in TEA at room temperature for antibody cross-linking. The beads were incubated for 15 min with 50 mM Tris (pH 7.5) at room temperature and washed three times with 0.05% PBS-T before resuspension in solubilized PM fractions and incubation overnight at 4°C. The beads were washed three times with 0.05% PBS-T and eluted with soft elution buffer [0.2% SDS (w/v), 0.1% Tween-20 (v/v), 50 mM Tris-HCl, pH 8.0] for BN-PAGE as outlined previously (Antrobus and Borner, 2011; Smalley et al., 2020).

### Densitometry

For Western blot analysis, bands from raw images were analyzed using the Image J densitometry features. Biological replicates were run on the same gels for comparison, and each band’s area under the curve was calculated. The average signal for each treatment group was calculated based on the protein expression levels.

### EEG surgeries and recording

Adult mice (8-9 weeks old) were subjected to EEG surgeries as described in Colmers et al., 2024. To implant EEG head mounts, the mice were given anesthesia using isoflurane inhalation, and 3-channel EEG/EMG head mounts (Pinnacle Technology) were superglued to the skull in alignment with lambda. For cortical recordings, four screws were placed above the frontal and parietal lobes, and silver epoxy was applied to each screw head to improve electrical connectivity between the electrodes and the head mount. After the surgery, the mice recovered for seven days in their home cages before experimentation. On the day of recording, the mice were connected to the pre-amps in the recording chambers for an hour to get accustomed to the environmental conditions before recording. First, a two-hour-long baseline recording was obtained; subsequently, the mice received a 20 mg/kg kainate injection (*i.p*.). Two hours later, the mice were administered with a single dose of 5 mg/kg Diazepam (*i.p*.) and were recorded for another hour to assess the effectiveness of OV350 treatment during the postnatal development period in mitigating kainate-induced *status epilepticus* (SE) in adult *Cdkl5* KO mice.

### EEG analysis

To assess the potential impact of OV350 treatment on baseline EEG power, a 20-minute silent period was analyzed, during which no muscular movement was detected based on the EMG channel. Mice received a single dose of 50 mg/kg OV350 or vehicle control for 12 consecutive days during the postnatal development period from postnatal day 10 to 21. To evaluate the effectiveness of OV350 in restoring the efficacy of diazepam, a 40-minute EEG signal was analyzed and compared to the vehicle group.

The EEG signals were transformed into the frequency domain to create power spectral plots using the fast-Fourier transformation (FFT). These plots were generated with LabChart software, employing an 8K FFT size, a Hann (cosine-bell) window, and 87.5% window overlap. The EEG frequency analysis involved binning the total signal into various frequency bands: delta (0-4 Hz), theta (4-8 Hz), alpha (8-13 Hz), beta (13-30 Hz), and gamma (30-100 Hz). The contribution of each frequency band to the total power was subsequently determined and compared across the treatment groups.

To identify and score seizures and SE, EEG recordings were examined to determine the onset of the first seizure and instances of SE. Seizure activity was characterized as any event exceeding 2.5 times the standard deviation of the preceding one minute of EEG activity, persisting for a minimum of 20 seconds. Status epilepticus was defined as seizure activity extending beyond 30 minutes or as continuous events separated by less than 30 seconds that returned to baseline (Arshad and Naegele, 2020). To investigate the effectiveness of OV350 in preventing the onset of diazepam-resistant SE, we compared the EEG epochs of 40 minutes following diazepam administration across all three groups of mice.

### Behavior

For all behavioral tests, wild-type mice aged seven to eight weeks were administered a vehicle only, while CDD mice were given injections of either a vehicle or OV350. The mice were acclimatized in the testing rooms for one hour before the tests. The tests were conducted at similar times each day (between 8 AM and 1 PM) in temperature-controlled rooms (24–26°C). Littermates were consistently tested at the same time. After each experiment, the mice were returned to their original cage. The equipment was sanitized after each mouse using 70% ethanol, followed by Clidox (a chlorine dioxide-based sterilant). Both male and female mice were utilized for all experiments.

### Barnes Maze Assay

Mice aged between 10 and 12 weeks were used for the assay. The maze used was a circular platform with a 1.5 m diameter, containing 40 holes along the perimeter, each with a 2.5 cm diameter. An escape tunnel was positioned under one of these holes. Mice used natural spatial cues in the room to locate the escape hole under the maze. The test procedure involved placing the mice in the center of the maze under a wire cage, allowing them to become familiar with the environment. Afterward, the cage was lifted, and the mice were given 3 minutes to explore the maze and find the escape hole. The time taken to exit the maze through the escape tunnel was then recorded. If the mice couldn’t find the escape within 3 minutes, their time was noted as the maximum of 180 seconds, and they were removed from the maze. This test was repeated thrice daily with a 30-minute interval between trials. The average duration for each day was considered for analysis. This protocol was followed for four consecutive days.

To assess short-term memory, the escape hole was removed on the fifth day, and the mice were given 5 minutes to explore the arena. The time spent at each hole was measured and put into 45° bins, with each containing five holes. These bins were organized around the perimeter of the maze in a clockwise direction, with 0° representing the goal hole and the adjacent holes. The same assessment was repeated on day 12 after a week of no exposure to the maze for long-term memory. An overhead camera and Ethovision software were used to track the time spent in each area of the arena.

### 3-Chamber Social Interaction Assay

At 13 weeks of age, mice were tested in a 3-chamber setup, with each chamber measuring 40 cm × 40 cm. The mice were placed in the center chamber and given 10 minutes to explore all the chambers. Metal cages measuring 4 cm × 4 cm × 5 cm with 1 cm gaps between each vertical bar were then placed in the center of the left and right chambers. The test mice were allowed to explore the arena for 5 minutes for habituation. After this, an unfamiliar male or female mouse (8 weeks old) was placed under one of the cages, and a dummy mouse was placed under the second cage, and the test mice were allowed to explore the arena for 10 minutes. The time spent in the chamber with the mouse was compared between all the mice. The time spent in the chamber with the familiar versus the dummy mouse was also calculated. An overhead camera and Ethovision software were used to detect time spent in each region of the arena.

### Statistical Analysis

All data are presented as the mean ± standard error of the mean (SEM). Shapiro-Wilk test was performed to find the normal distribution of the data sets. Biochemistry data were analyzed using the Mann-Whitney’s test or t -test. EEG data was analyzed using the Two-Way ANOVA followed by Tukey’s Multiple Comparison. Behavioral data were analyzed using the Two-Way ANOVA to compare between genotypes followed by Tukey’s Multiple Comparison, and the Repeated Measures ANOVA followed by Dunnet’s Multiple Comparison to analyze data obtained on individual mice across different trials. Data are shown in Table 2 (including p-values, test used, means, standard error of the mean, and the number of animals). P values < 0.05 are considered statistically significant.

**Table 2.**
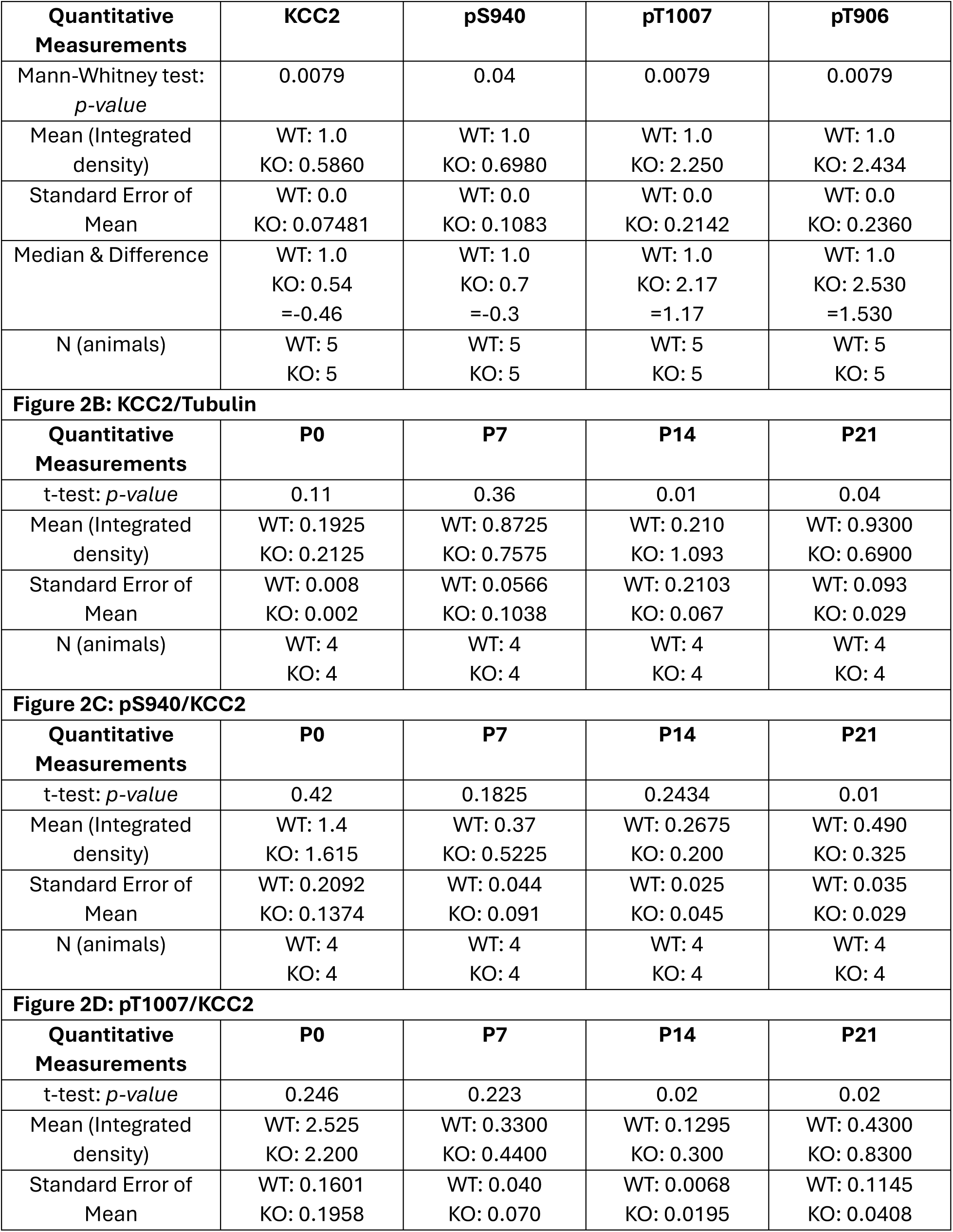
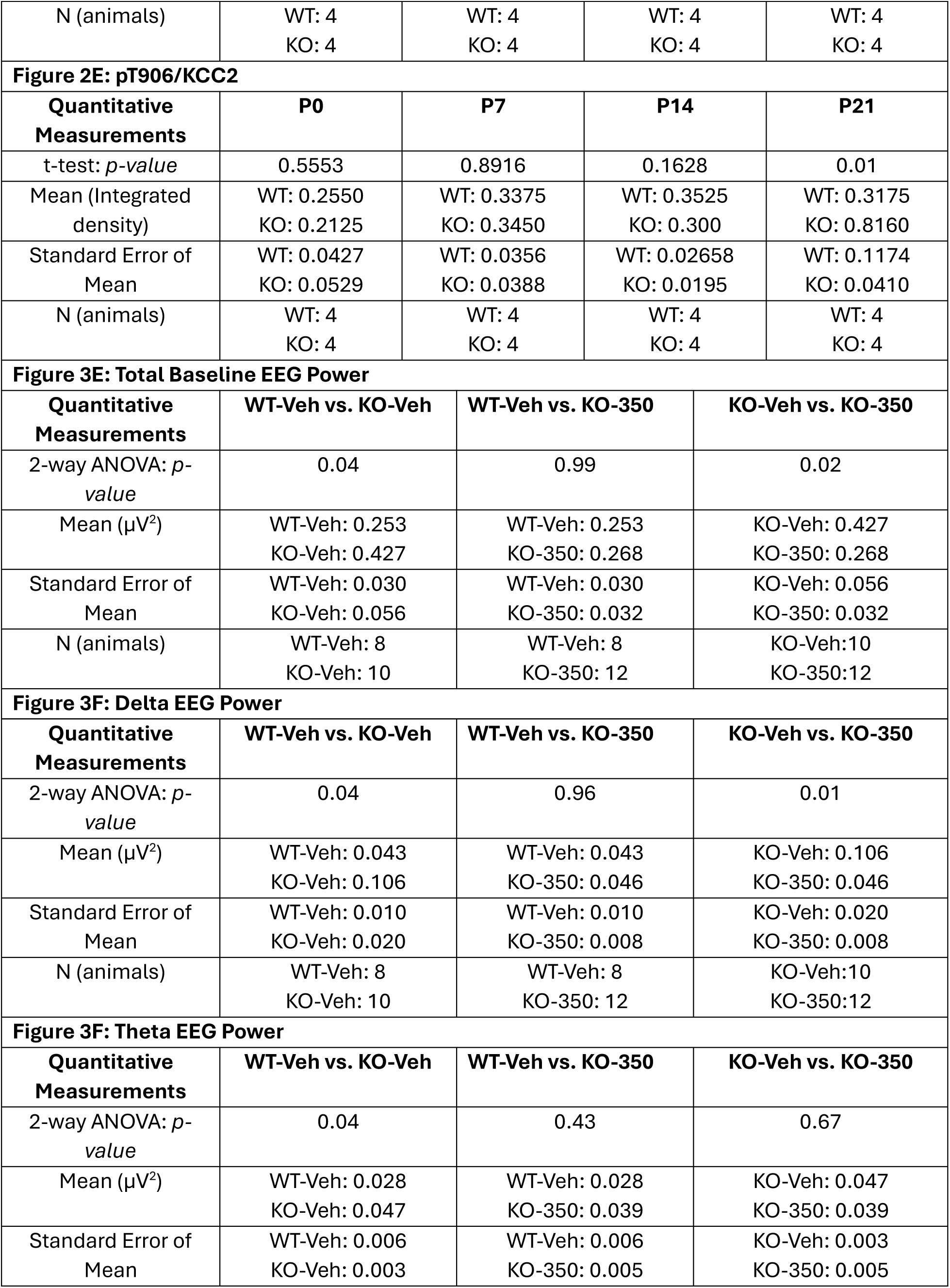
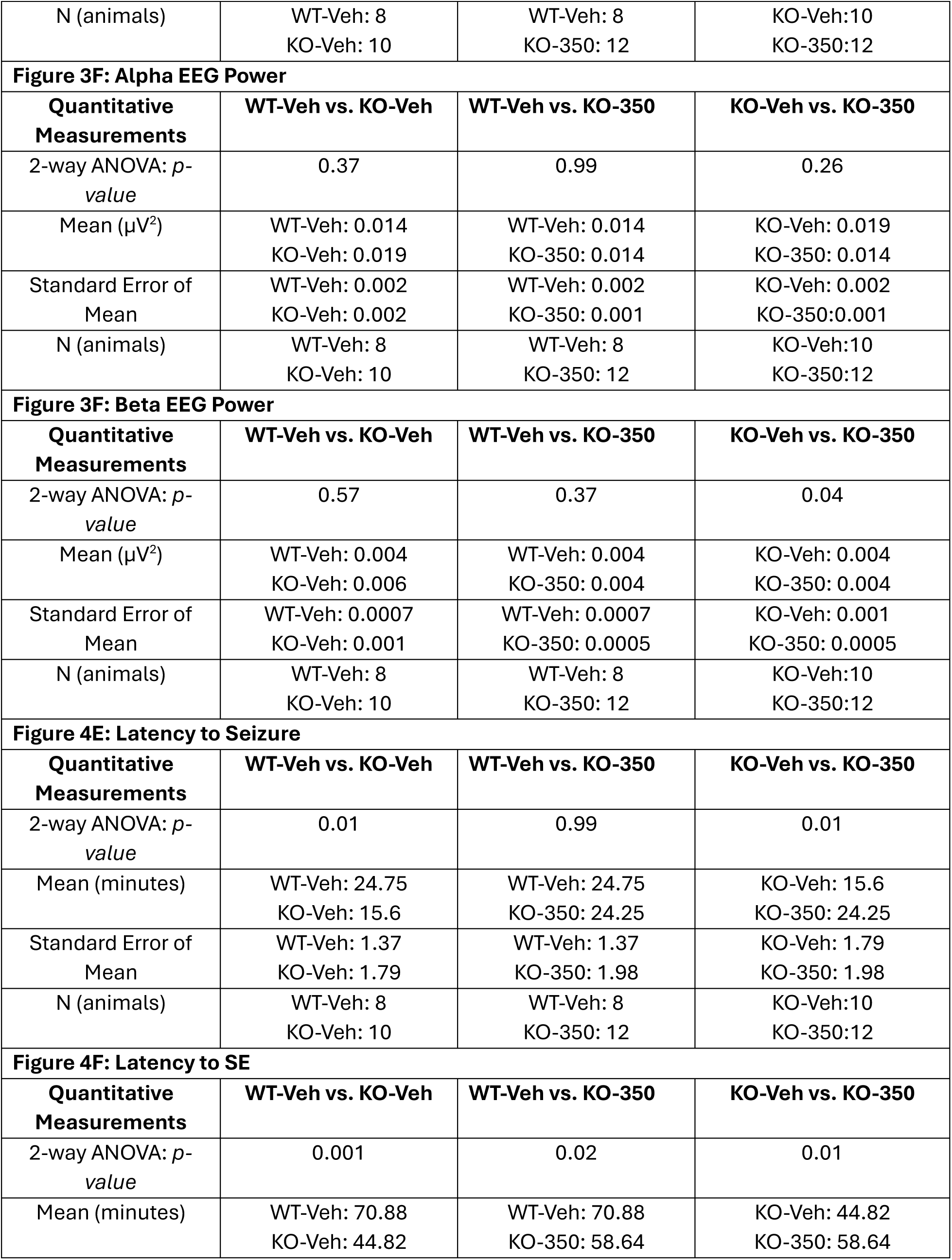
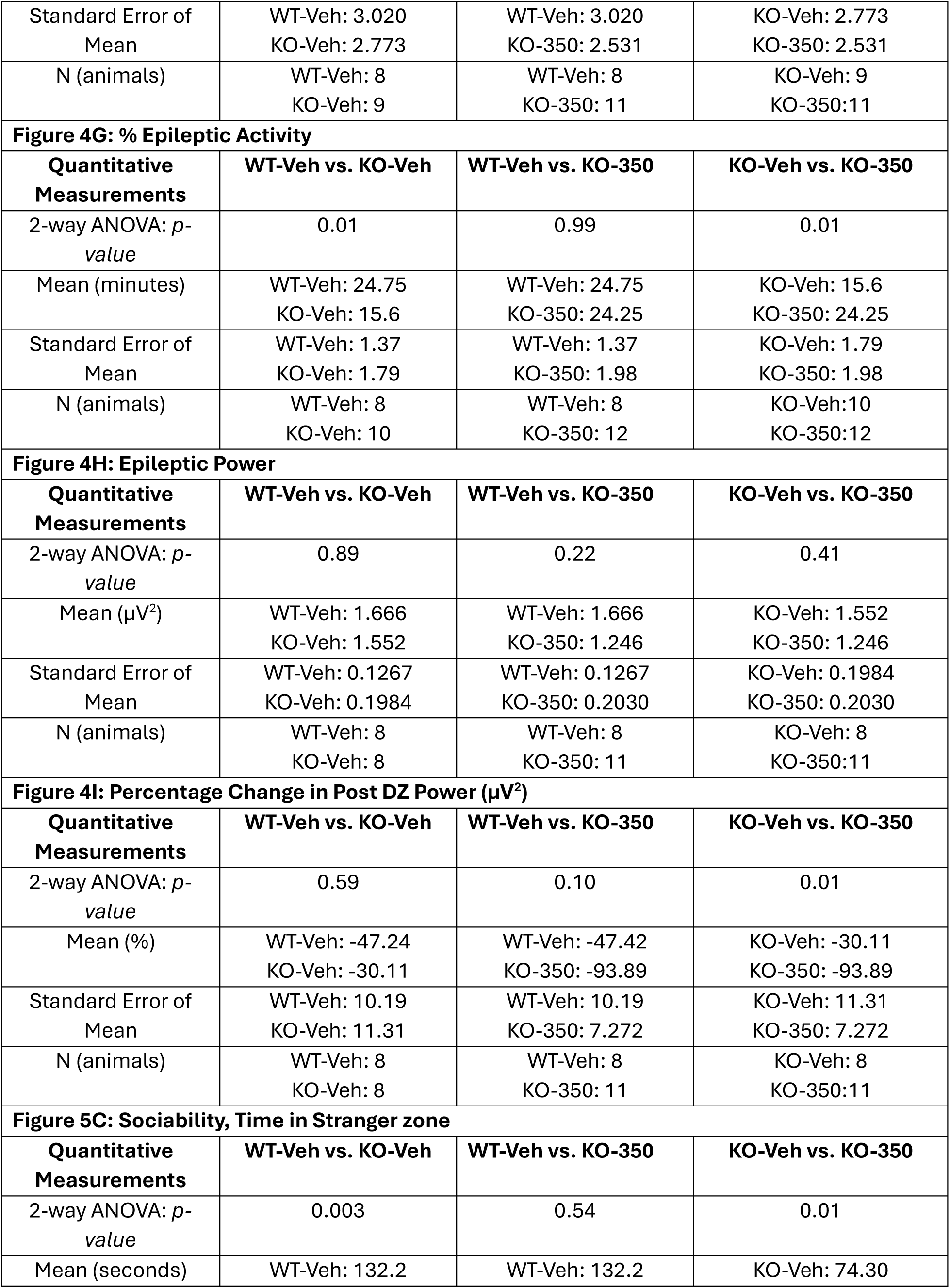
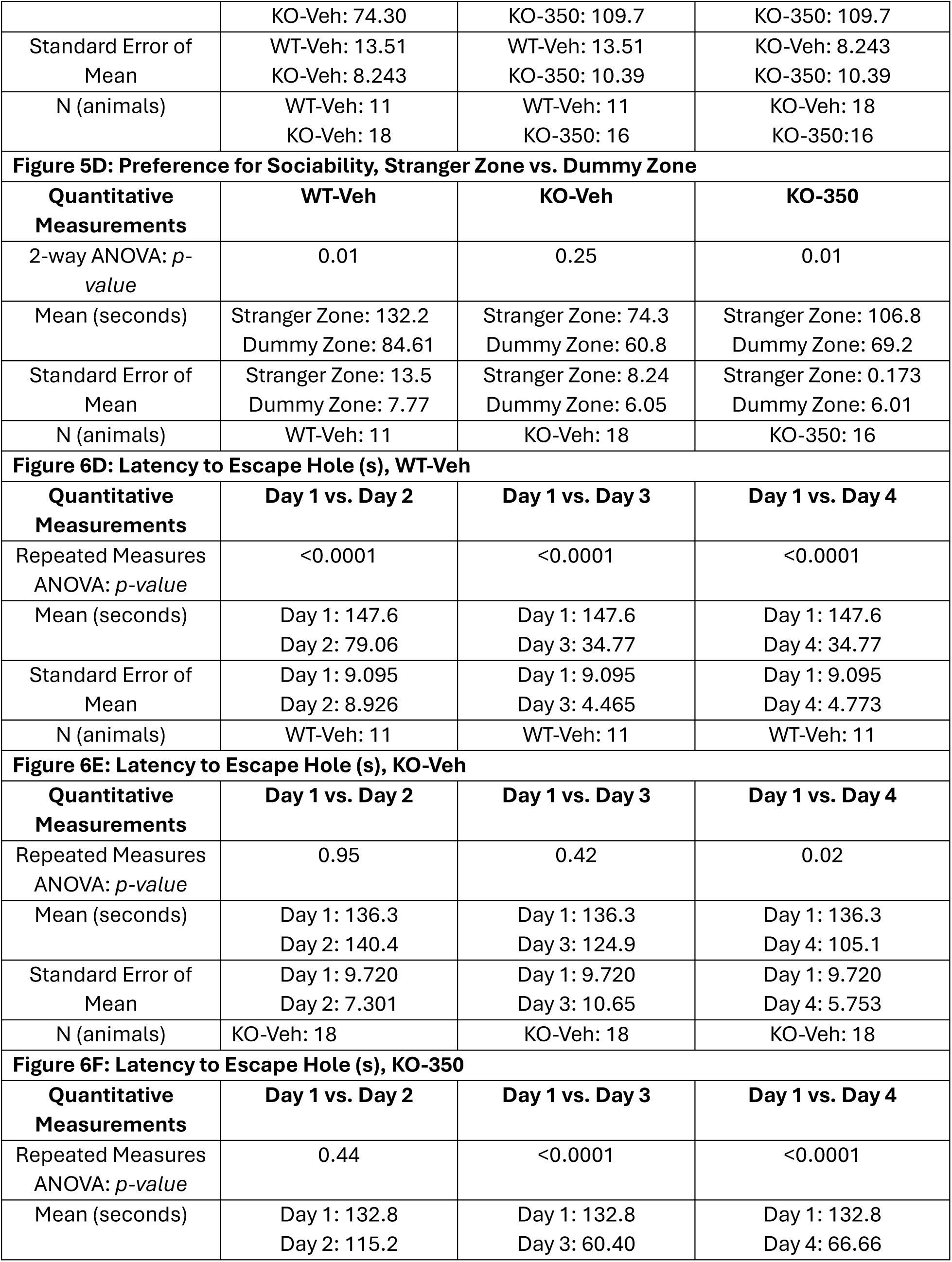
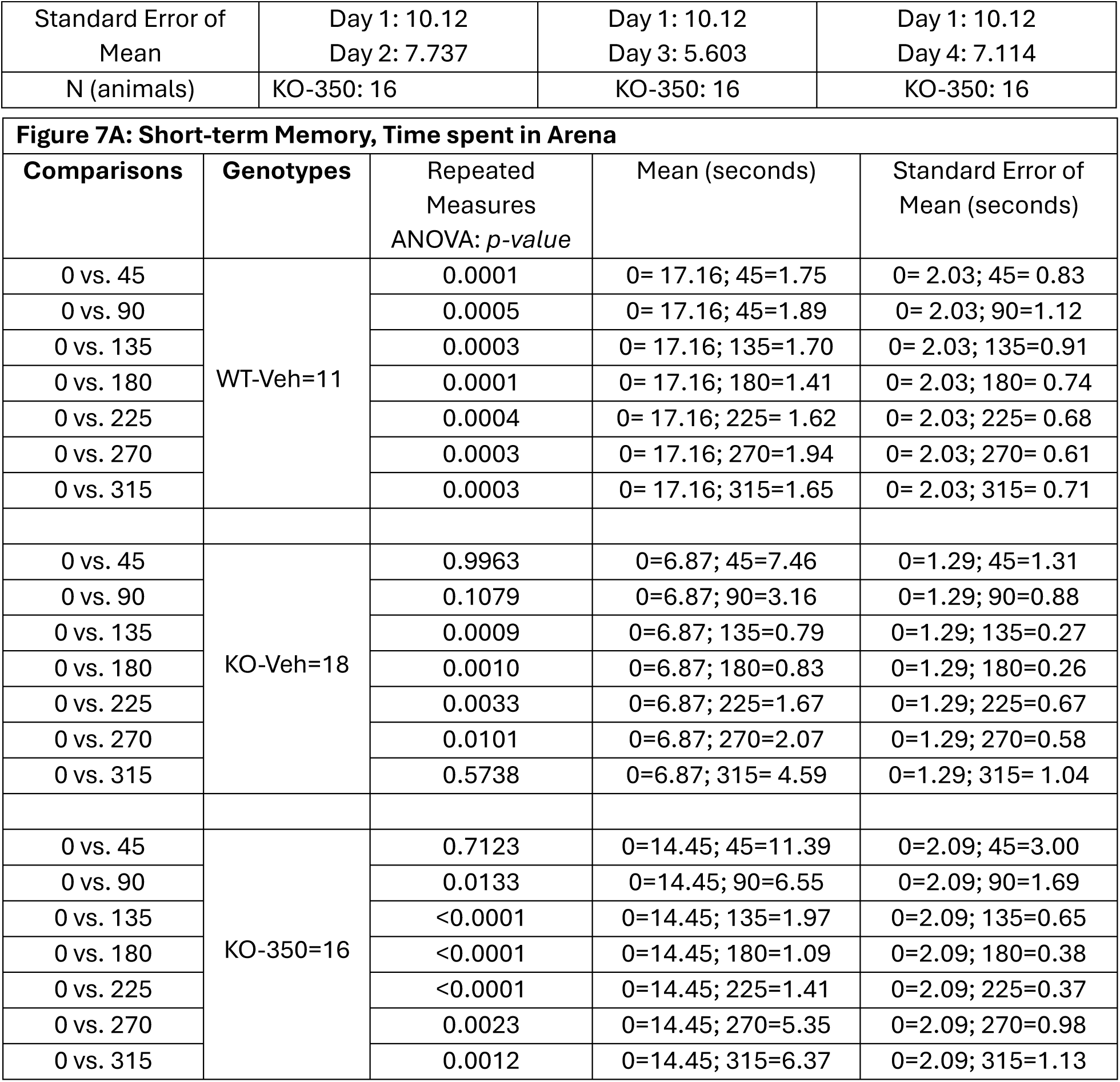
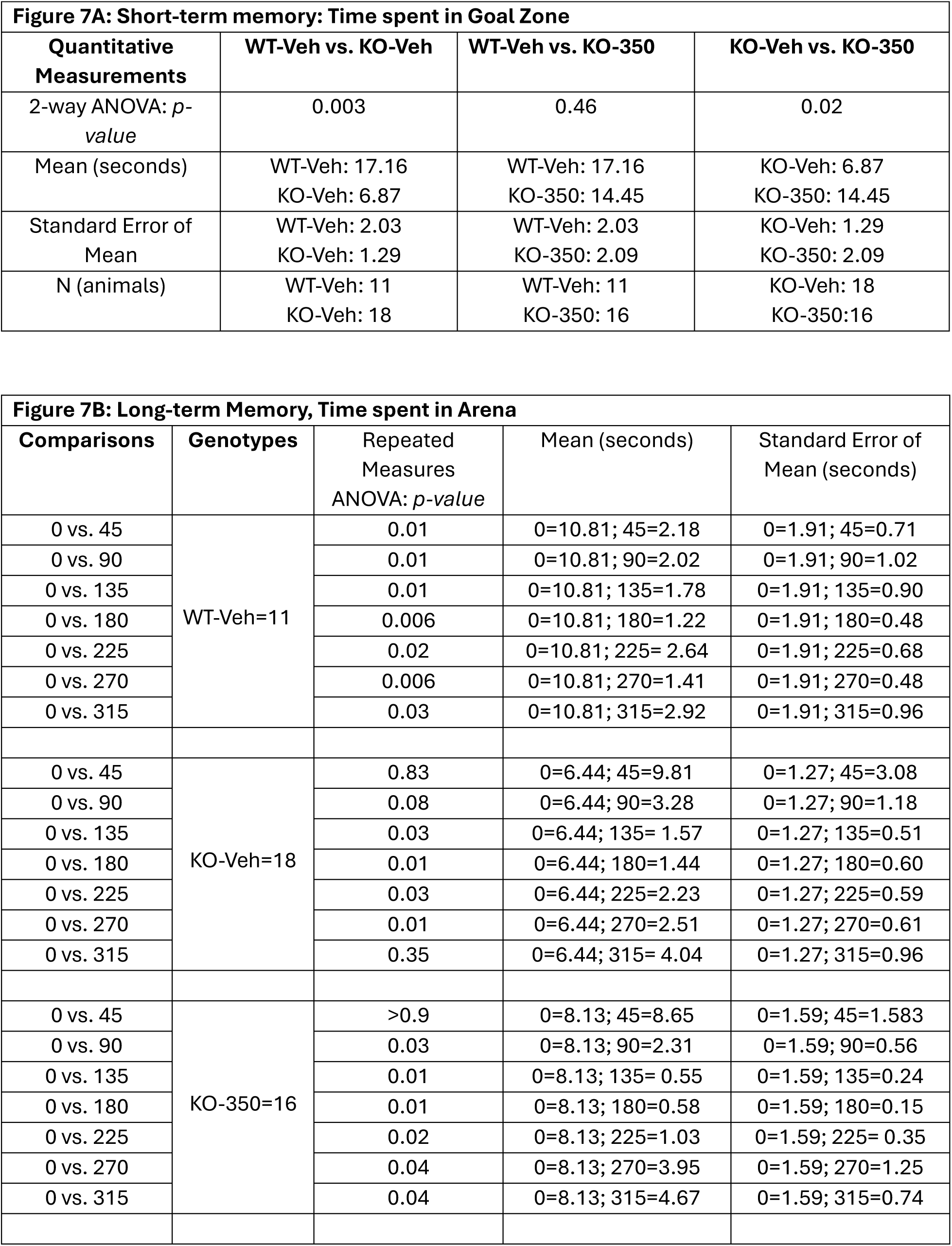
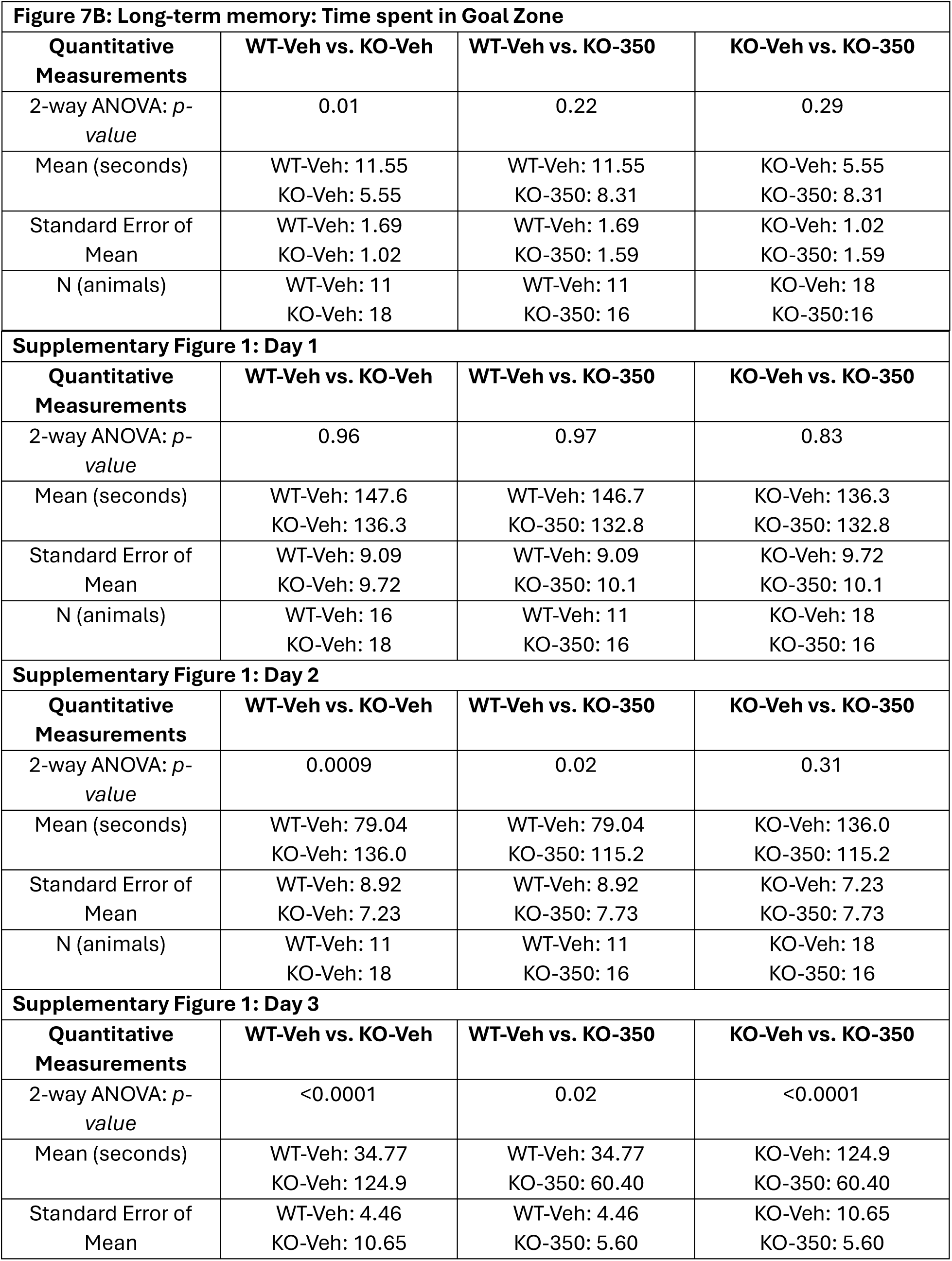
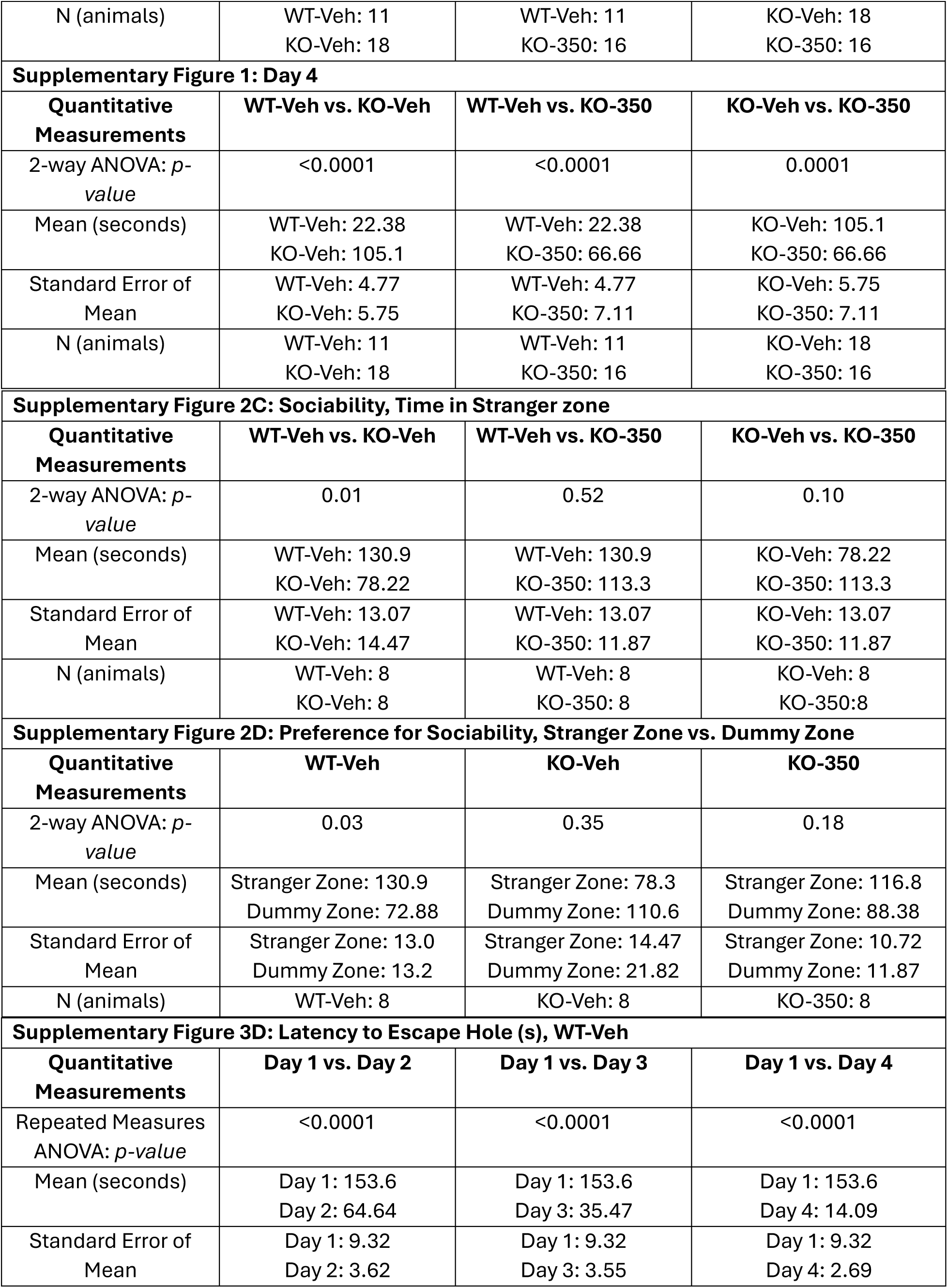
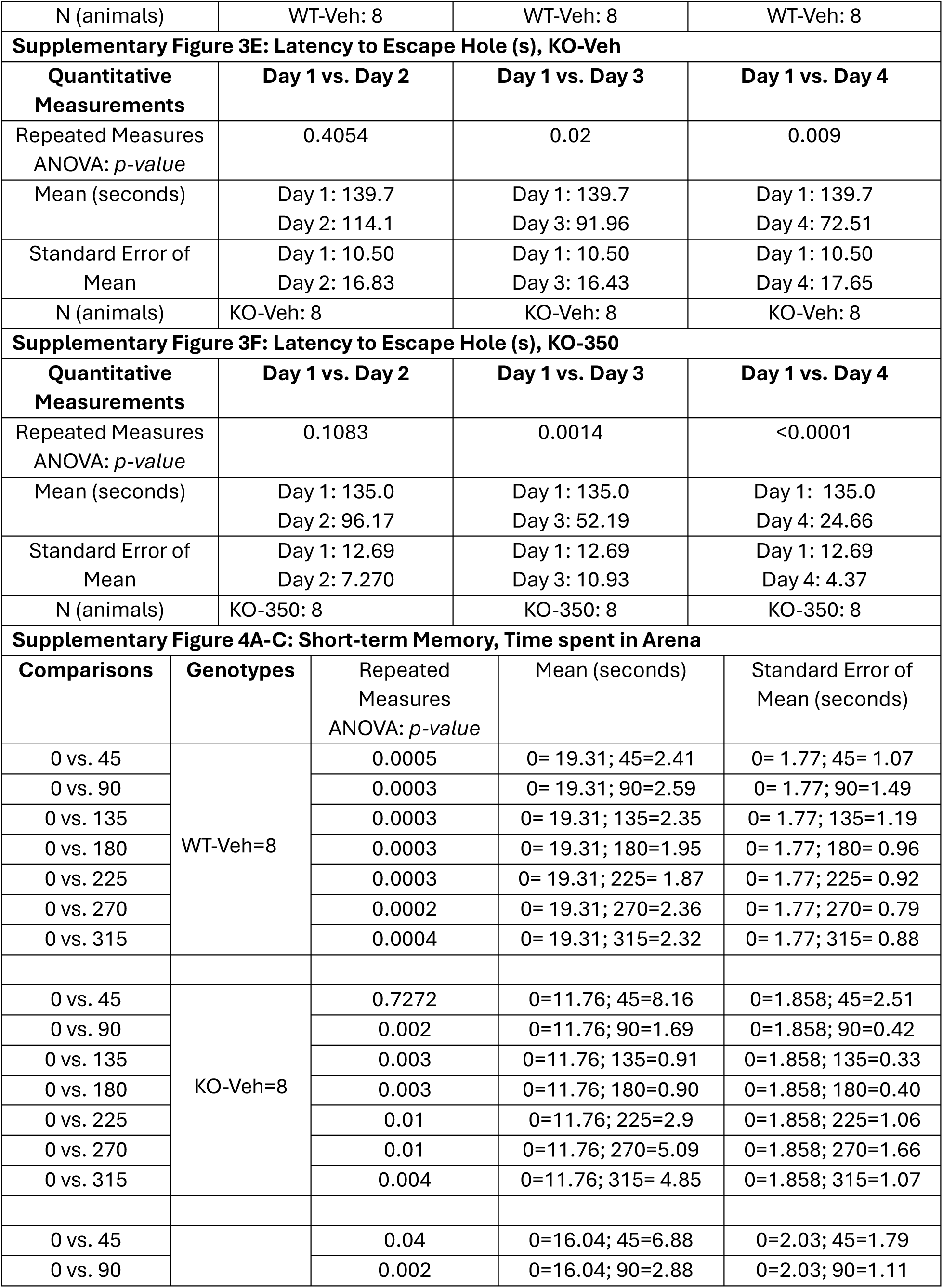
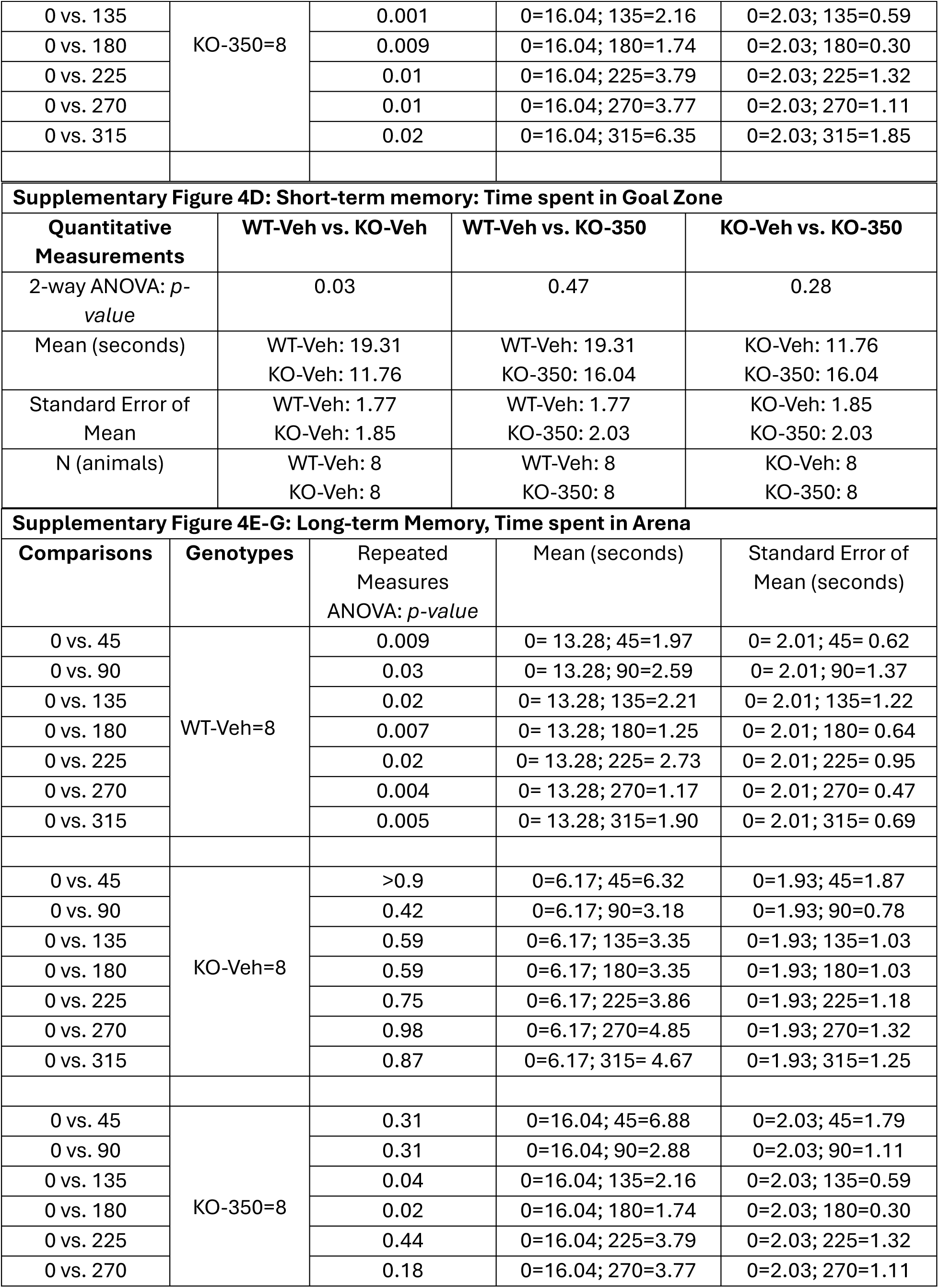
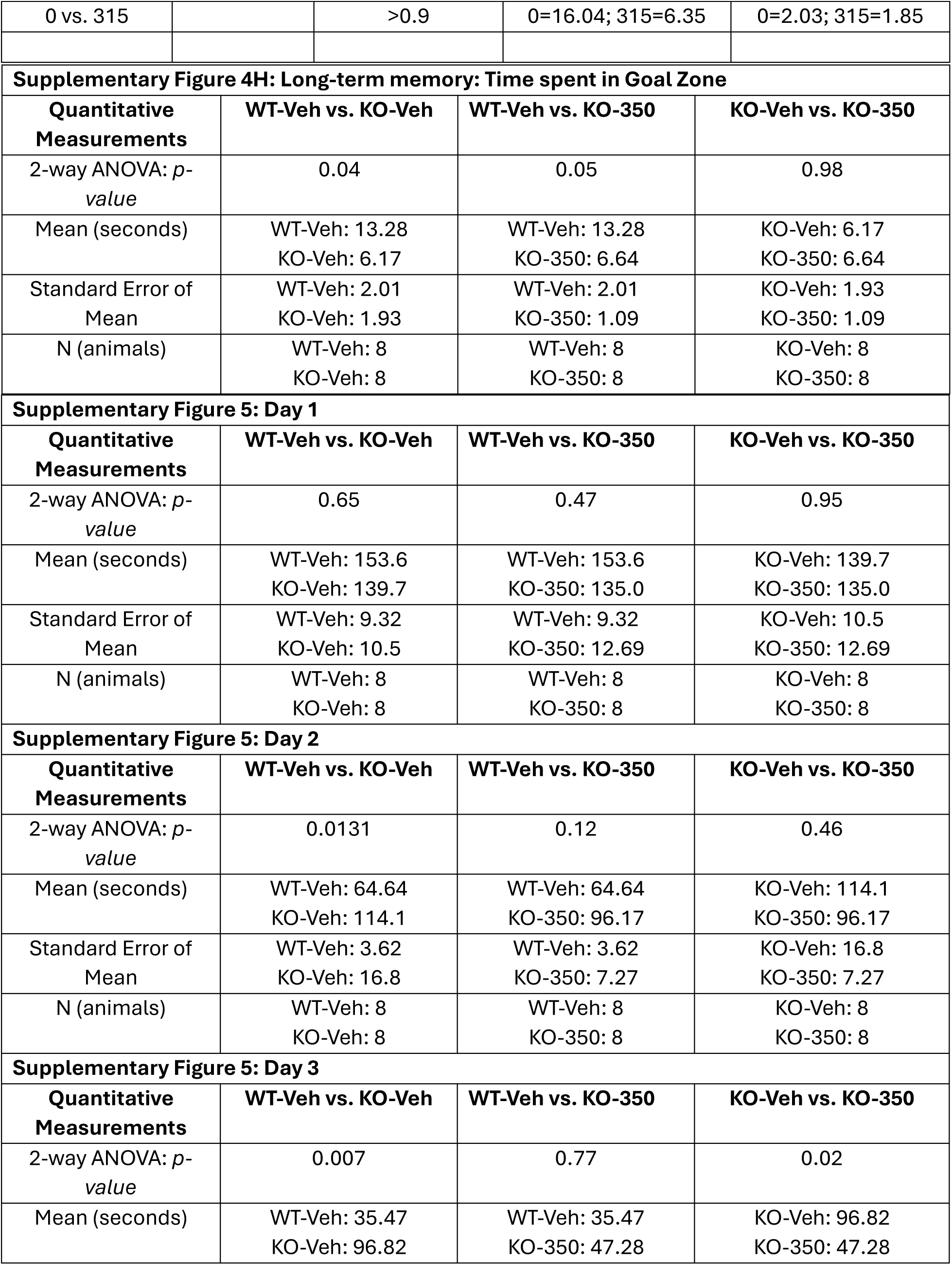
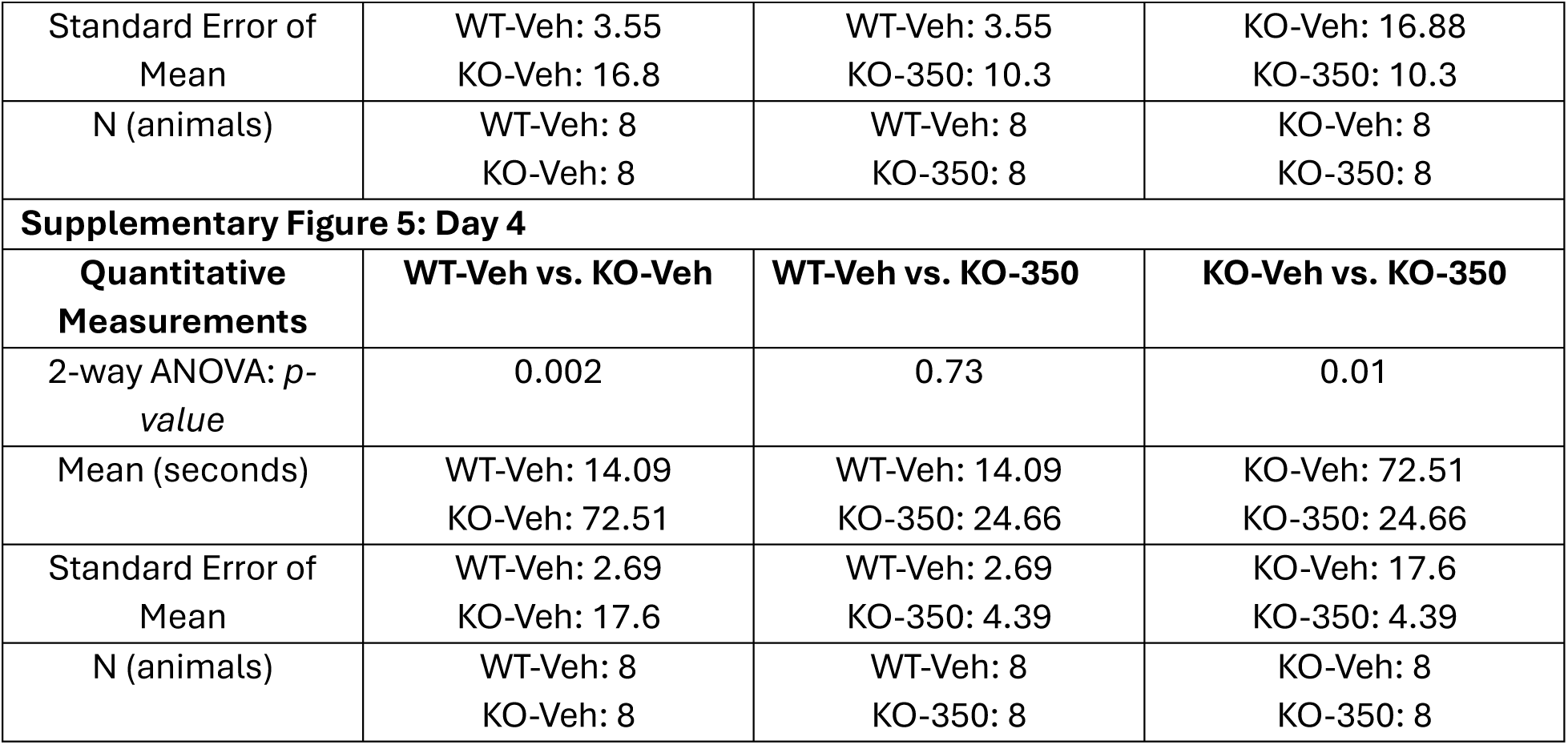
Statistical Analysis.

## Results

### Loss of functional CDKL5 is associated with aberrant KCC2 phosphorylation and expression

To confirm the association between CDKL5 and KCC2, we isolated the cortex and hippocampus and made purified plasma membrane preparations that were subjected to immunoprecipitation with KCC2 or control IgG antibodies. Precipitated material was immunoblotted with CDKL5 antibody. In WT mice, CDKL5 is associated with KCC2 and is part of the KCC2 complex. CDKL5 was significantly reduced in the *Cdkl5* KO mice (Figure 1A). Cortical and hippocampal lysates prepared from WT or *Cdkl5* KO mice (8-12 weeks old) were immunoblotted with antibodies against β-actin, CDKL5, KCC2, and phospho-specific antibodies pS940, pT906, and pT1007. Compared to WT mice, there was a significant reduction in KCC2 expression in *Cdkl5* KO mice (Figure 1B-C). There was a concomitant reduction in the phosphorylation level of KCC2 at the serine 940 site (Figure 1B, 1D). In contrast, phosphorylation is increased at T906 (Figure 1B, 1E) and T1007 (Figure 1B, 1F) sites in *Cdkl5* KO mice.

**Figure 1.**
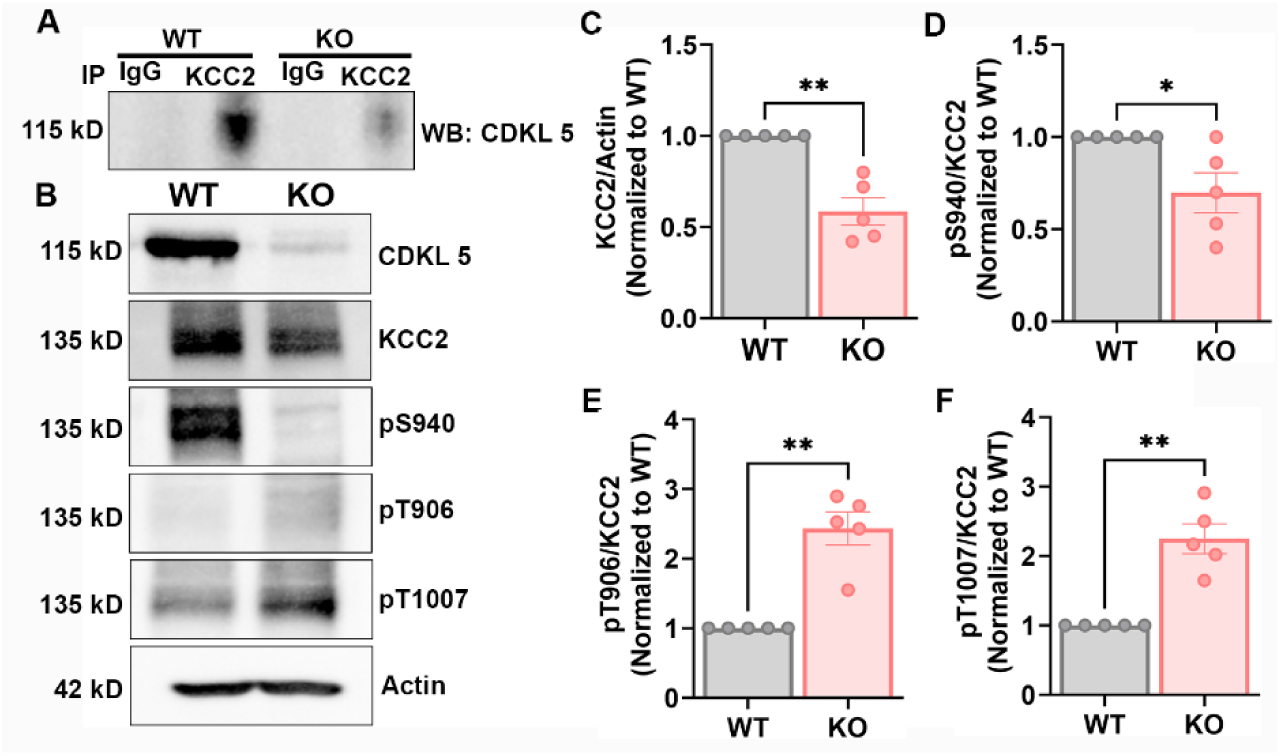
CDKL5 is strongly associated with the KCC2 complex, and loss of *Cdkl5* modifies KCC2 phosphorylation. **A**) Immunoprecipitation of KCC2 complex in plasma membrane preparations shows a strong association of CDKL5 with KCC2 in the WT mice when blotted with Cdkl5 antibody. CDKL5 is reduced in the KCC2 complex in *Cdkl5* KO mice. **B**) Forebrain lysates prepared from the WT, or *Cdkl5* KO mice were immunoblotted with beta-actin, Cdkl5, KCC2, and phospho-specific antibodies against S940(pS940), T906(pT906), and T1007(pT1007). **C**) Graph shows a reduction in KCC2 expression in *Cdkl5* KO mice compared to the WT mice. **D**) S940 phosphorylation of KCC2 is reduced in *Cdkl5* KO mice compared to the WT mice. **E**) T906 phosphorylation of KCC2 is increased in the *Cdkl5* KO mice compared to the WT mice. **F**) KCC2 phosphorylation at T1007 is increased in *Cdkl5* KO mice compared to the WT mice. The asterisk indicates the significance of the *Cdkl5* KO mouse compared to WT. See table 2 for statistical analysis.

Thus, these results demonstrate that loss of CDKL5 results in a KCC2 phosphorylation phenotype that displays the hallmarks of an immature phenotype. We have previously published that such phosphorylation changes lead to decreased KCC2 function, which is strongly correlated with excessive neuronal excitation, increased seizure susceptibility, and cognitive, and behavioral impairments in adult mice (Silayeva et al., 2015; Moore et al., 2018; Moore et al., 2019).

### CDKL5 loss orchestrates alteration in postnatal regulation of KCC2 expression and phosphorylation

Phosphorylation of KCC2 residue serine 940 (S940), threonine 1007, and threonine 906 regulate KCC2 function in the adult brain, so we sought to examine the developmental profile of S940, T906, and T1007 phosphorylation. We measured total KCC2 expression and KCC2 S940, T906, and T1007 phosphorylation in whole brain lysates at p0, p7, p14, and p21 and detected phosphorylation of this residue at each of these time points (Figure 2A). Total KCC2 expression increased linearly over-development till p14, followed by maintenance at this level as the brain neurons further matured by p21 in WT mice, with significantly lower expression at p14 and p21 days in *Cdkl5* KO pups compared to the WT pups (Figure 2B). We detected a significant decrease in S940 phosphorylation at p21 between WT and *Cdkl5* KO pups (Figure 2C). Phosphorylation of KCC2 at threonine residues 906 and 1007 decreases during development in WT mice, which contributes to the developmental upregulation of KCC2 function. Therefore, we also compared the phosphorylation status at these residues between WT and *Cdkl5* KO pups. We observed significant increases in KCC2 phosphorylation at T1007 from p14 onwards in *Cdkl5* KO pups compared to WT pups (Figure 2D). We also detected a significant increase in phosphorylation at T906 at p21 age in *Cdkl5* KO pups compared to the WT pups (Figure 2E). An increase in phosphorylation in these threonine residues during development in *Cdkl5* KO pups suggests a developmental delay in the maturation of KCC2.

**Figure 2.**
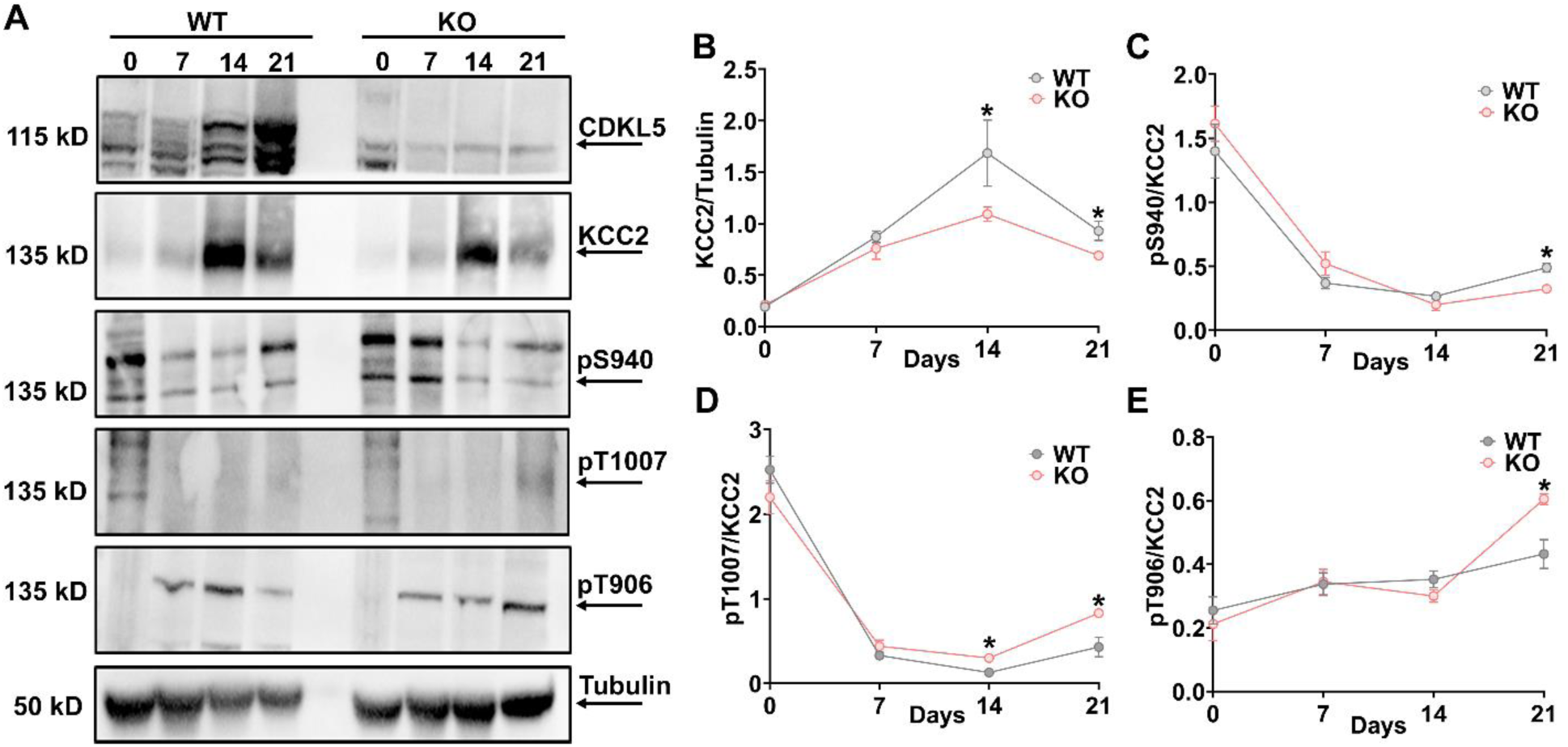
CDKL5 ablation alters KCC2 expression and phosphorylation over time during postnatal development. **A**) We performed western blotting to assess developmental changes in KCC2 expression and phosphorylation in WT and Cdkl5 KO pups at 0, 7, 14, and 21 postnatal days. Whole brain lysates prepared from WT, or Cdkl5 KO pups were immunoblotted with beta-tubulin, CDKL5, KCC2, and phospho-specific antibodies against S940(pS940), T906(pT906), and T1007(pT1007). **B**) KCC2 expression progressively increased from p0 to p14 and stabilized by p21 in WT and *Cdkl5* KO pups. However, KCC2 expression from p14 to p21 is significantly reduced in *Cdkl5* KO mice compared to the WT pups. **C**) Relative to levels of KCC2 expression, S940 phosphorylation is decreased from p0 to p21 in WT and *Cdkl5* KO pups. The S940 phosphorylation was significantly reduced at p21 in *Cdkl5* KO pups compared to the WT pups. **D**) Graph shows a progressive reduction in T1007 phosphorylation over time in WT pups relative to the KCC2 expression. Meanwhile, *Cdkl5* ablation significantly increased phosphorylation at the T1007 site on KCC2 from p14 to p21 compared to WT pups. **E**) The T906 phosphorylation on KCC2 doesn’t change over time from p0-p21 during postnatal development in WT and *Cdkl5* KO mice. However, the phosphorylation is significantly increased at the T906 site on KCC2 at p21 in *Cdkl5 KO* pups. The asterisk indicates the significance of the *Cdkl5* KO mouse compared to WT on a particular day. See table 2 for statistical analysis.

### Adult *Cdkl5* KO mice exhibit increased baseline EEG power, which is normalized by OV350 treatment during the postnatal developmental period

The postnatal change in KCC2 phosphorylation regulates the developmental GABA switch. Disruption to the timing of the switch (as would be expected in *Cdkl5* KO mice with an immature KCC2 phosphorylation signature) will have long lasting consequences for many developing brain circuits, altering their long-term excitability (Furukawa et al., 2017; Reh et al., 2021). Here, we used EEG recordings to measure baseline EEG signal in adult WT and *Cdkl5* KO mice. We further examined the consequences of enhancing KCC2 activity during this critical developmental period on baseline EEG measurements. To address this hypothesis, we administered a daily dose of the KCC2 activator (OV350, 50 mg/kg *i.p*.) to *Cdkl5* KO pups from p10-p21, compared to vehicle treated, age-matched, WT and *Cdkl5* KO pups. After four weeks, EEG surgeries were performed. In the following week, after recovering from the surgeries, a two-hour-long baseline recording was conducted to examine the difference in baseline EEG power between the WT-Veh, *Cdkl5* KO-Veh, and *Cdkl5* KO-OV350 mice (Figure 3A). A 20-minute-long EEG epoch recording was compared between all three groups. To quantify the difference, recordings were subjected to fast Fourier transformation (FFT) to convert the EEG signals from the time domain into the frequency domain, generating a power spectral density plot for frequencies between 0 and 100 Hz (Figure 3B-D). The total baseline EEG power of *Cdkl5* KO-Veh mice was significantly higher than the WT mice (Figure 3E). Treatment of infant *Cdkl5* KO mice with OV350 between p10 and p21 resulted in their adult EEG baseline power being significantly reduced compared to vehicle-treated *Cdkl5* KO mice, with power being comparable to the WT mice (Figure 3E). We next assessed the possible differences in the EEG frequency bands (delta; 0–4 Hz, theta; 4–8 Hz, alpha; 8–13 Hz, and beta; 13–30 Hz) and observed a significant increase in power in the delta and theta frequency bands (Figure 3F). However, in adult *Cdkl5* KO mice treated with OV350 while they were infants, the delta and theta frequency bands were not significantly different to WT vehicle-treated mice (Figure 3F). These results suggest that the *Cdkl5* KO mice may be more prone to seizures due to the higher baseline EEG power. Therefore, we compared the susceptibility to seizures and *status epilepticus* (SE) between the two groups of mice.

**Figure 3.**
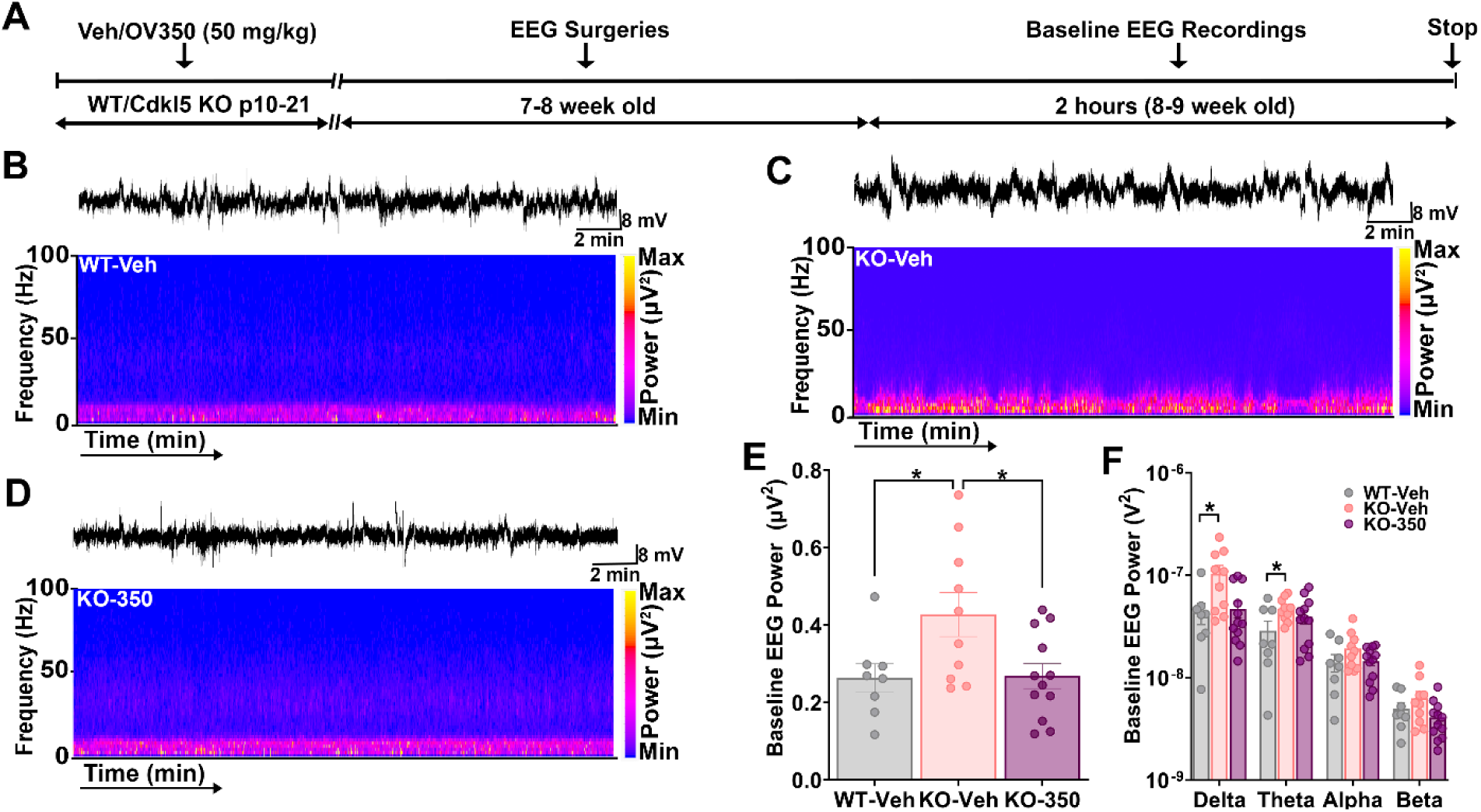
KCC2 activator reduces the baseline EEG power in *Cdkl5* KO mice. **A)** The line diagram shows an experimental timeline. Two hours of baseline EEG recording was performed, followed by vehicle/OV350 (50mg/kg i.p., arrow) administration, and recordings were performed for an additional two hours. **B)** Representative EEG trace and its spectrogram show power distribution across different frequency bands from a vehicle-treated wild-type mouse. **C)** Representative EEG trace is shown above power spectra from a vehicle-treated *Cdkl5* KO mouse. **D**) Representative EEG trace is shown above power spectra from a vehicle-treated *Cdkl5* KO mouse. **E)** Baseline EEG recordings from all three groups of mice were subjected to FFT, and a spectral plot is shown for frequencies between 0-100 Hz. Total EEG power was compared between the three groups of mice. *Cdkl5* KO mice had a higher baseline EEG power than WT mice. Mice treated with OV350 significantly reduced the EEG power compared to the vehicle-treated *Cdkl5* KO mice. The asterisk indicates the significant difference in baseline EEG power among different groups of mice. **F)** The OV350 treatment did not change the basal EEG power across all frequency bands. The asterisk indicates the significant difference in power of a particular frequency band among different groups of mice. See table 2 for statistical analysis.

### *Cdkl5* KO mice are more susceptible to kainate-induced seizures and status epilepticus

Since KCC2 phosphorylation influences seizure severity, *Cdkl5* KO mice with altered KCC2 phosphorylation would be predicted to have increased duration of epileptiform activity, a decreased latency to the first epileptiform event and a decreased latency to onset of SE compared to WT mice. To address this hypothesis, we compared the susceptibility of kainate-induced seizures in wild-type and *Cdkl5* KO mice.. A single dose of 20 mg/kg kainate was administered, with recordings continuing for 2 hours. After that, all three groups of mice received a saturating concentration of diazepam (DZ; 5 mg/kg i.p.), after which recordings were extended for an additional 1 hour (Figure 4A-D).

**Figure 4.**
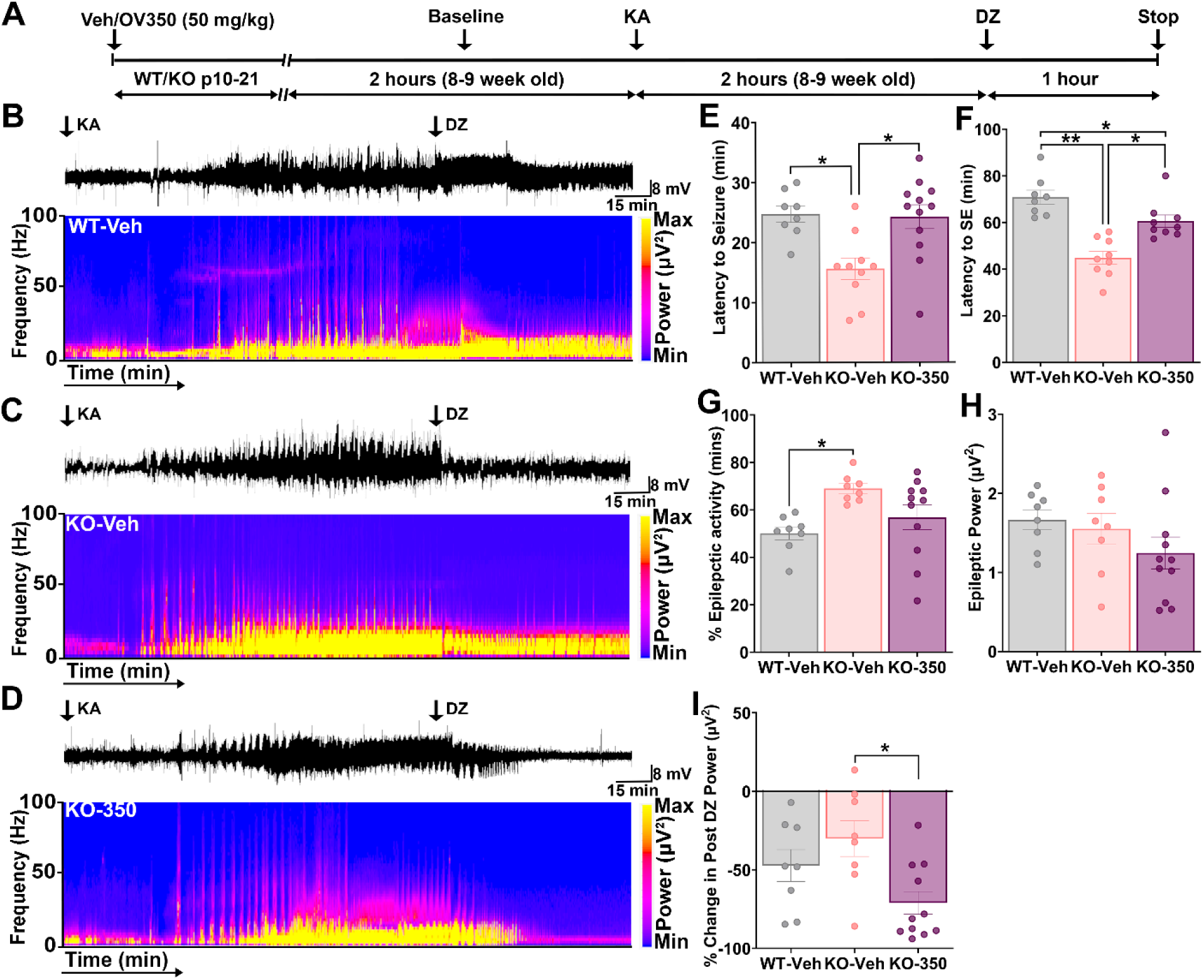
KCC2 activator increases the susceptibility to kainate-induced seizures and SE in *Cdkl5* KO mice. **A)** Experimental timeline shows that wild-type pups were administered with the vehicle, and *Cdkl5* KO pups received either vehicle or OV350 (50 mg/kg *i.p*.) from p10-p21 during the postnatal development period. Four weeks later, these mice underwent EEG surgeries. One week following the surgeries, on the day of the experiment, a 2 hour of baseline recording was conducted. Next, mice received a single injection of kainate (20mg/kg i.p., first arrow) to induce seizures and status epilepticus. Two hours after KA injection, mice were dosed (i.p.) with 5 mg/kg DZ, and EEG recordings were extended for 1 hour. **B)** Representative EEG trace and its spectrogram show power distribution across different frequency bands from a vehicle-treated WT mouse. **C)** Representative EEG trace and spectrogram show power distribution across different frequency bands from a vehicle-treated *Cdkl5* KO mouse. **D)** A representative EEG trace is shown above in the power spectra of an OV350-treated *Cdkl5* KO mouse. **E)** Graph shows increased latency to the first seizure in *Cdkl5* KO mice. The asterisk indicates the significant difference in latency to the first seizure among different groups of mice. **F)** OV350-treated *Cdkl5* KO mice took more time to get into KA- induced SE. The asterisk indicates the significant difference in latency to the SE among different groups of mice. **G**) Vehicle-treated wildtype mice spent less time in epileptic events than the vehicle-treated *Cdkl5* KO mice, and OV350 treatment did not reduce time spent in epileptic activity in the *Cdkl5* KO mice. The asterisk indicates the significant difference in time spent in epileptic activity among different groups of mice. **H)** EEG recordings of the epileptic activity from the vehicle and OV350 treated mice were subjected to FFT, and spectral plot is shown for frequencies between 0-100 Hz. Total EEG power was compared between all three groups of mice. OV350 treatment did not alter the EEG power in *Cdkl5* KO mice. **I)** EEG recordings of the post-diazepam treatment period from all three groups of mice were subjected to FFT, and a spectral plot is shown for frequencies between 0-100 Hz. Total EEG power was compared. Mice treated with OV350 significantly reduced the EEG power compared to the vehicle-treated mice. The asterisk indicates the significant difference in change in post-DZ power among different groups of mice. See table 2 for statistical analysis.

This model was chosen because of the similarities with patients having drug-resistant seizures, as KA-induced seizures become refractory to benzodiazepines within minutes. The *Cdkl5* KO mice took less time to have their first seizure (Figure 4E) and developed SE faster than WT mice (Figure 4F). Additionally, the *Cdkl5* KO mice spent more time in epileptic activity than the WT mice (Figure 4G). However, we did not observe a difference in the EEG power of total epileptic activity (Figure 4H). In both WT and *Cdkl5* KO mice, the benzodiazepine, diazepam, failed to suppress SE, as demonstrated by the lack of change in EEG power prior to and post-diazepam treatment (Figure 4I).

### Targeting KCC2 activity in infant *Cdkl5* KO mice reduces seizure susceptibility, and restores diazepam efficacy, in adult mice

We next explored if these seizure susceptibility issues observed in adult *Cdkl5* KO mice could be negated by enhancing KCC2 function during development. Adult *Cdkl5* KO mice that received OV350 treatment between p10-p21 had a significantly increased latency to the first kainate-induced seizure compared with vehicle-treated *Cdkl5* KO mice (Figure 4E). These mice also took a significantly longer time to develop status epilepticus than the vehicle-treated *Cdkl5* KO mice (Figure 4F). However, KCC2 activation didn’t reduce the time spent in epileptic activity and epileptic power in *Cdkl5* KO mice (Figure 4G-H).

Consistent with previously published studies using the KA model, DZ did not modify EEG power in mice treated with a vehicle. In contrast, in *Cdkl5* KO mice treated with OV350 (50 mg/kg *i.p*.) between p10-p21, DZ significantly reduced EEG power in adult mice (Figure 4I).

### OV350 treatment during postnatal development improves sociability in adult *Cdkl5* KO mice

The *Cdkl5* KO mice exhibit autistic-like behavioral abnormalities and perform poorly in social interaction tasks (Jhang et al., 2017; Fuchs et al., 2018). Deficits in KCC2 expression and phosphorylation also promote autism-like behavior in mice (Pisella et al., 2019; Virtanen et al., 2021). However, increasing KCC2 activity during development improves sociability in mice (Moore et al, 2019). Therefore, we investigated the impact of enhancing KCC2 activity using OV350 (50 mg/kg *i.p*.) during development (p10-21) on social behavior in adult *Cdkl5* KO mice (10-11 weeks) (Figure 5A). We hypothesized that the abnormal social behavior in *Cdkl5* KO mice is linked to altered KCC2 phosphorylation and reduced KCC2 activity, and potentiating KCC2 activity during development would rescue sociability in *Cdkl5* KO mice. Sociability was assessed using a 3-chamber social interaction test (Figure 5B). Firstly, sociability was examined by measuring time spent interacting with an unfamiliar (stranger) mouse. Consistent with the literature *Cdkl5* KO-Veh mice spent significantly less time interacting with the stranger mouse than the WT-Veh mice did, indicating that the *Cdkl5* deficient mice have a reduced motivation for social interaction. However, the OV350-treated *Cdkl5* KO mice spent more time interacting with the stranger mouse compared to the time *Cdkl5* KO-Veh mice spent, suggesting that potentiating KCC2 activity during development alleviates deficits in sociability in *Cdkl5* deficient mice (Figure 5C). Next, we measured how much time mice spend with a live mouse compared to a dummy mouse. *Cdkl5* KO-Veh treated mice spent almost the same time with live and dummy mice. However, *Cdkl5* KO-OV350 mice, like WT mice, spent more time with a live mouse (Figure 5D). This finding suggests that KCC2 activation during development in *Cdkl5* KO mice increases their preference for sociability as adult mice.

**Figure 5.**
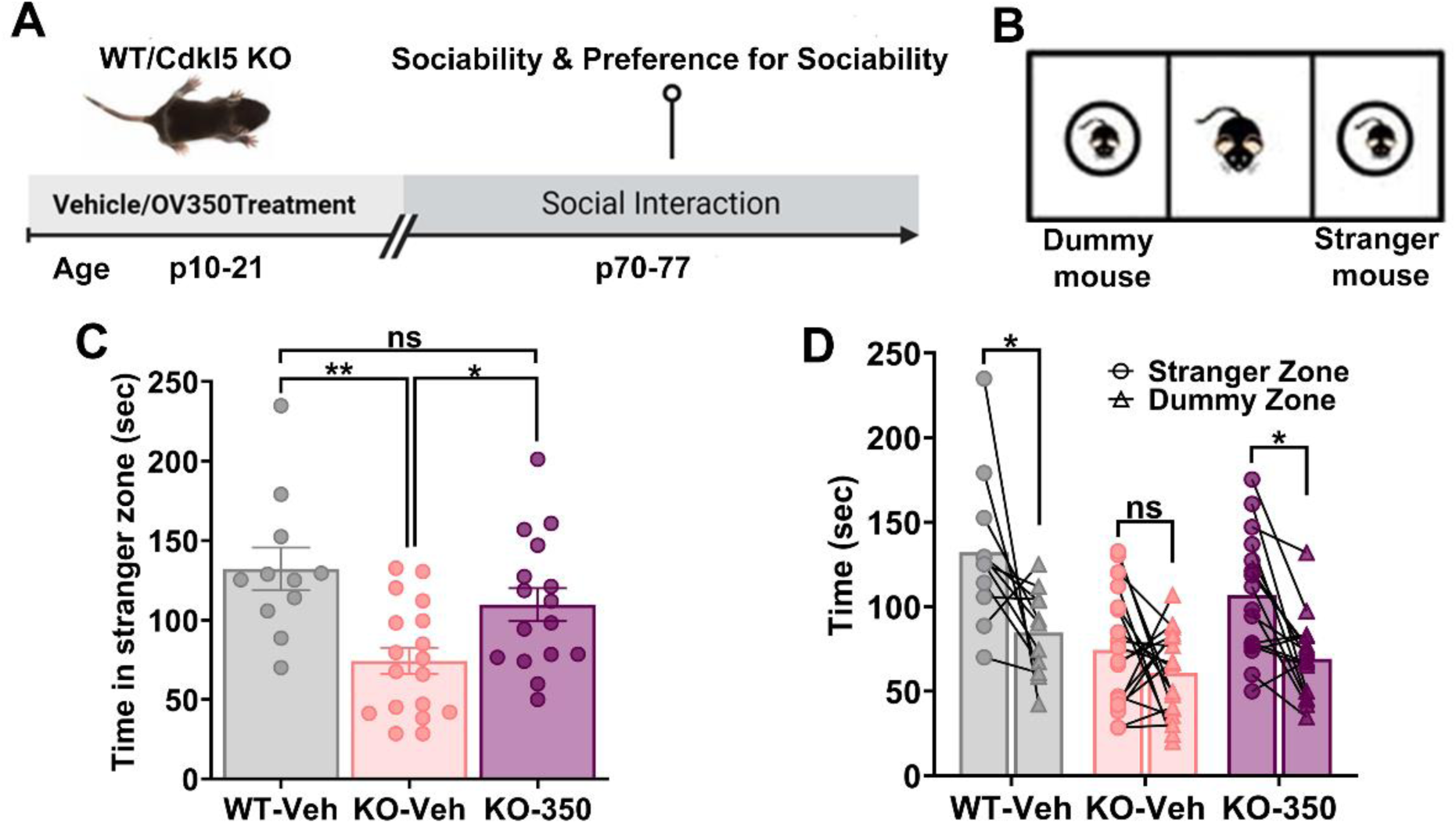
KCC2 activation during development improves sociability in *Cdkl5* KO mice. **A)** Experimental timeline. The WT pups were administered a single dose of vehicle daily from p10-p21. The *Cdkl5* KO pups received a single dose of either vehicle or OV350 (50 mg/kg *i.p*.) daily from p10-p21. We used a three-chamber social interaction assay to assess sociability and preference for social familiarity in the *Cdkl5* KO mice treated with or without OV350. **B)** Diagram of the sociability assay. Mice were allowed to explore either an unfamiliar mouse or a dummy mouse. **C)** Wild-type mice spent more time interacting with the stranger mouse than vehicle-treated *Cdkl5* KO mice, and OV350-treated *Cdkl5* KO mice spent more time interacting with the stranger mouse than vehicle-treated mouse. The asterisk indicates the significance of the time spent with the stranger mouse. **D)** Graph shows that both WT and OV350-treated *Cdkl5 KO* mice preferred interacting with the mouse vs. the dummy mouse. The asterisk indicates the significance of the time spent with the dummy and the stranger mouse. See table 2 for statistical analysis.

### Administering KCC2 activator during postnatal development improves spatial learning in *Cdkl5* KO mice

Intellectual disabilities are a core feature of CDD, and compromised KCC2 phosphorylation also impacts learning and memory. We performed a Barnes maze assay to examine spatial learning and memory in the *Cdkl5* KO mice. We treated *Cdkl5* KO mice with OV350 (50 mg/kg *i.p*.) or Vehicle during development (p10-21), and wild-type mice received vehicle injections only. We then examined the beneficial effects of increasing KCC2 activity during development on spatial memory in adult *Cdkl5* KO mice (8-10 weeks old) (Figure 6A-B). To assess the impact of treatment on learning, we recorded the latencies to enter the escape hole over 4 days of learning (Figure 6C). *Cdkl5* KO-Veh mice performed poorly in escape latencies compared to the wildtype mice over the 4-day learning period, while the OV350-treated mice performed better than the vehicle-treated *Cdkl5* KO mice but not as well as the wildtype mice (Supplementary figure 1). The wild-type mice showed a significant improvement over their day one escape latencies by day two (Figure 6D). The *Cdkl5* KO-vehicle showed an improvement in escape latencies by training day 4 (Figure 6E). In contrast, the *Cdkl5* KO-350 mice showed a significant improvement in escape latencies by training day 3 (Figure 6F). This suggests that the rate of spatial learning was mildly improved in the OV350-treated *Cdkl5* KO mice.

**Figure 6.**
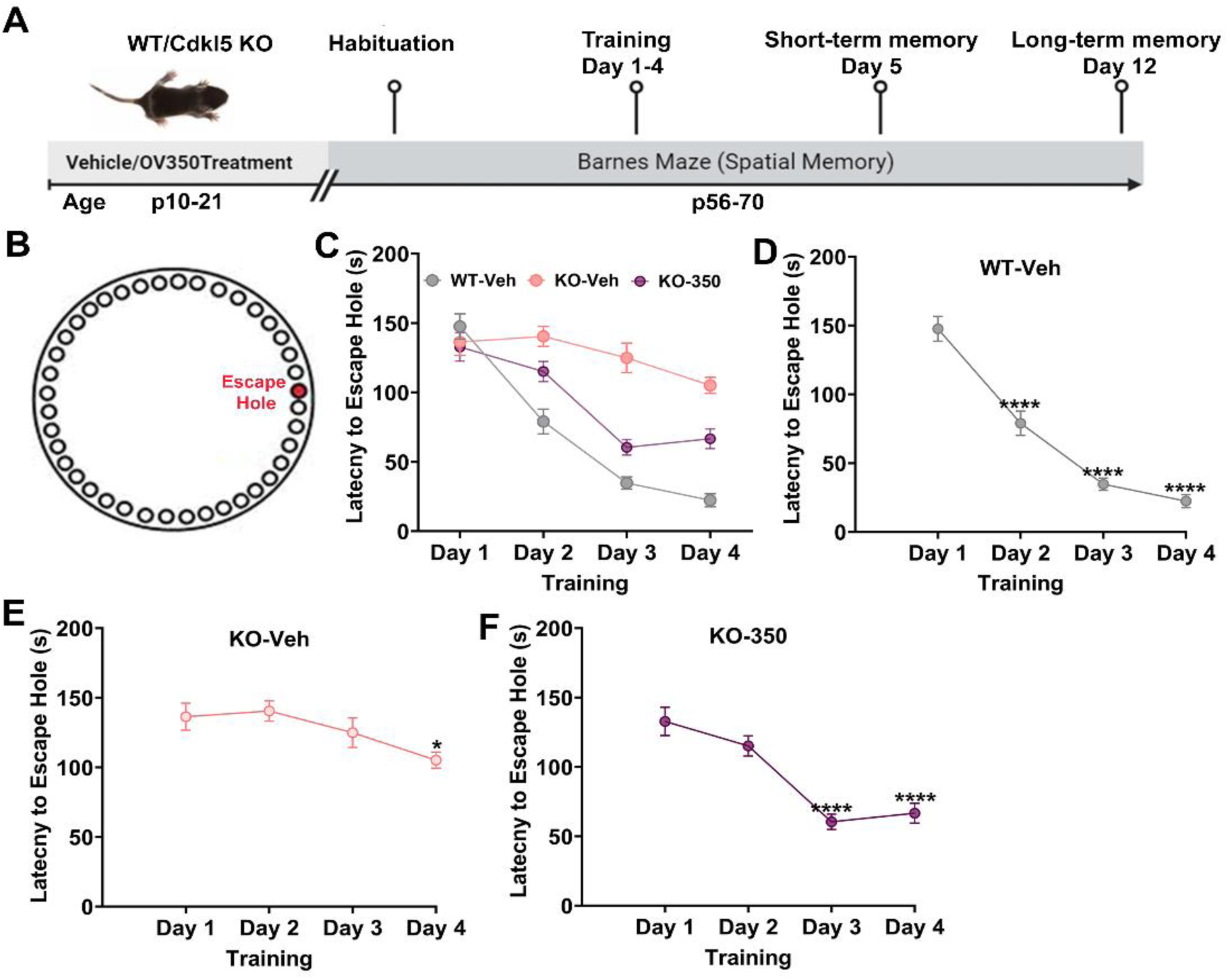
KCC2 activation during the postnatal development period improves the rate of spatial learning in *Cdkl5* KO mice. **A)** Experimental timeline. The *Cdkl5* KO pups received a single dose of either vehicle or OV350 (50 mg/kg *i.p*.) daily from p10-p21. WT pups were administered a single dose of vehicle daily from p10-p21. We used a Barnes maze assay to assess the rate of spatial learning in the WT-vehicle and *Cdkl5* KO mice treated with or without OV350. **B)** Cartoon illustration of the Barnes maze. **C)** Latency to enter the goal hole was measured on days 1–4; wild-type mice learned better than the vehicle-treated *Cdkl5* KO mice from days 2-4. However, OV350-treated *Cdkl5* KO mice performed considerably better than vehicle-treated *Cdkl5* KO mice from days 3-4. **D)** The learning rate (day when there is a significant reduction in latency to goal compared to day 1) was also comparable in wild-type mice. The asterisk indicates the significant difference in latency to the escape hole with reference to day 1. **E)** The learning rate was comparable in vehicle-treated *Cdkl5* KO mice from day 4. The asterisk indicates the significant difference in latency to the escape hole with reference to day 1. **F)** The learning rate was comparable in OV350-treated *Cdkl5* KO mice from day 3. The asterisk indicates the significant difference in latency to the escape hole with reference to day 1. See table 2 for statistical analysis.

### Potentiating KCC2 activity during postnatal development impacts spatial memory in *Cdkl5* KO mice

Memory assessments were conducted over increasing periods after the learning portion of the Barnes maze assay. We assessed short-term and long-term memory on day 5 and day 12, respectively. To examine memory retention, we removed the escape chamber to measure the time spent in the space where the escape hole was initially located. Time spent at the escape hole differed between wild-type and *Cdkl5* KO-vehicle mice on day 5, suggesting that the short-term memory was significantly impaired in the *Cdkl5* KO-vehicle mice. Interestingly, OV350-treated *Cdkl5* KO mice spent more time investigating the escape hole region than other regions in the maze on day 5 (Figure 7A). By day 12, the *Cdkl5* KO-vehicle mice couldn’t differentiate between the escape and non-escape regions, but the wild-type mice showed long-term memory retention and spent more time in the escape hole region. While the OV350-treated *Cdkl5* KO mice demonstrated a preference for the escape hole region compared to the non-escape hole region, it didn’t spend a significant time at the escape hole (Figure 7B), suggesting that increasing KCC2 function during development improves short-term spatial memory retention as adults, but has limited impact on long-term memory.

**Figure 7.**
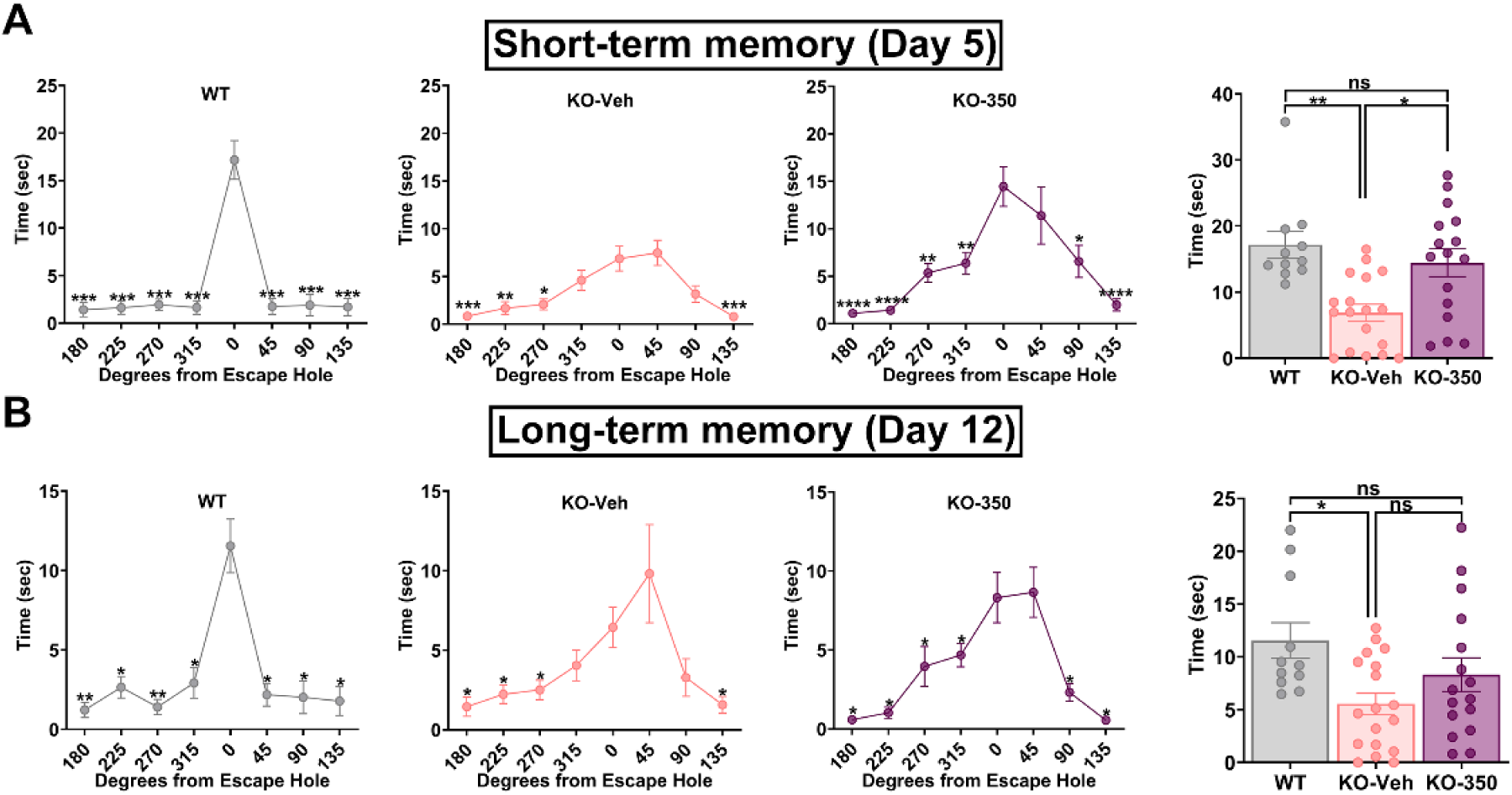
KCC2 activation during development improves spatial memory retention in *Cdkl5* KO mice. Spatial memory was assessed using a Barnes maze assay. After 4 days of training, time spent at each hole was measured on day 5 and day 12 upon removal of the escape tunnel, and data was binned into 45° groups. **A)** WT-vehicle mice spent more time in the goal area on day 5. Vehicle-treated *Cdkl5* KO mice didn’t perform as well as the WT mice. OV350-treated (50 mg/kg *i.p*.) *Cdkl5* KO mice performed considerably better. WT-vehicle and OV 350 treated *Cdkl5* KO mice spent more time at the goal zone than the vehicle-treated *Cdkl5* KO mice. B) WT mice show enhanced specificity of the spatial memory on day 12 compared to the vehicle and OV350-treated *Cdkl5* KO mice. The WT mice spent more time at the goal than the other two groups of mice. * Indicates significant difference to time spent in the 0° region. See table 2 for statistical analysis.

### OV350 treatment during adulthood doesn’t reduce sociability deficits in *Cdkl5* KO mice

Sociability and cognitive deficits persist into adulthood in both human patients and animal models of CDD. Therefore, we examined whether KCC2 activation during adulthood, when the neural circuits have already been established, could alleviate social deficits in *Cdkl5* KO mice (Supplementary Figure 2).

First, we compared the time spent with the stranger mouse between all three groups of mice. Consistent with the literature, we found that vehicle-treated *Cdkl5* KO mice spent significantly less time with the stranger mouse than wild-type mice did. We also found that the OV350 treatment (50 mg/kg *i.p*.) didn’t significantly improve the time spent with the stranger mouse compared to the vehicle-treated mice. However, the time spent with the stranger mouse was comparable between the wild type and *Cdkl5* KO-OV350 mice. Next, we compared the sociability of all three groups of mice with the live stranger mouse to the dummy mouse and found that wild-type mice preferred to spend time with the live mouse. However, the vehicle-treated *Cdkl5* KO mice could not differentiate between the live and the dummy mice. Similarly, the OV350-treated *Cdkl5 KO* mice didn’t spend significantly more time with the live mouse. This result suggests that contrary to the results found with OV350 treatment in infant *Cdkl5* KO mice, OV350 treatment in adults does not promote a preference for sociability in *Cdkl5* KO mice (Supplementary Figure 2).

### KCC2 activator treatment of *Cdkl5* KO mice during adulthood shows limited improvement in spatial learning and memory

We next sought to determine if *Cdkl5* KO mice treated as adults with OV350 can rescue the cognitive deficits. We used a Barnes maze assay to assess learning and memory, and latency to enter over four consecutive days (Supplementary Figure 3). Adult *Cdkl5* KO-Vehicle mice performed poorly in escape latencies compared to the wild-type mice over the 4-day learning period. In contrast, the OV350-treated (50 mg/kg *i.p*.) *Cdkl5* KO mice performed better than the vehicle-treated *Cdkl5 KO* mice, though still not as well as the wild-type mice. The *Cdkl5* KO-350 mice showed a significant improvement over their day one escape latencies by training day 3. This suggests that the rate of spatial learning was mildly improved in adult *Cdkl5* KO mice treated with OV350 (Supplementary Figure 3).

Memory assessments were conducted over increasing periods of time after the learning portion of the Barnes maze assay. We removed the escape chamber on days 5 and 12 and then measured the time spent in the space where the escape hole was originally located. Time spent at the goal hole differed between wild-type and *Cdkl5* KO-vehicle mice on day 5 and day 12 (Supplementary Figure 4). Memory was significantly impaired in the *Cdkl5* KO-vehicle mice on day 5. However, the performance of OV350-treated mice was not significantly different to that of wild-type mice and vehicle-treated *Cdkl5* KO mice. By day 12, *Cdkl5* KO-vehicle mice spent equal amounts of time at the non-goal regions as they did at the goal region, demonstrating a lack of spatial memory specificity compared to their wildtype mice, suggesting that *Cdkl5* KO-vehicle mice have a deficit in memory retention.

In contrast, *Cdkl5* KO-350 mice spent more time at the goal hole position on both day 5 and day 12 than the *Cdkl5* KO-vehicle mice, suggesting that increasing KCC2 function may improve memory retention (Supplementary Figure 4).

## Discussion

Our findings suggest that direct activation of KCC2 during postnatal development in *Cdkl5* KO mice normalize baseline EEG power and enhances the effectiveness of diazepam in terminating drug-induced status epilepticus (SE). Additionally, we observed that KCC2 activation during development alleviates sociability and cognitive deficits in *Cdkl5* KO mice.

KCC2 functions by extruding intracellular chloride anions and is essential for the ontogenetic switch of GABA_A_-mediated responses from depolarizing to hyperpolarizing (Virtanen et al., 2021). The expression and function of KCC2 play a crucial role in regulating neuronal excitability, and disruptions in KCC2 have been associated with the onset of infantile epilepsy (McMoneagle et al., 2024). Our findings indicate a significant decrease in KCC2 expression during postnatal development, as well as in adult *Cdkl5* KO mice. This reduction in KCC2 expression may result from the activation of the BDNF-TrkB pathway in the *Cdkl5* KO mice (Zhu et al., 2023). Notably, BDNF levels are elevated in *Cdkl5* KO mice, which is known to negatively impact KCC2 expression (Rivera et al., 2002). It is well established that phosphorylation of KCC2 at the serine 940 site (pS940) is critical for stabilizing KCC2 on the neuronal membrane surface, enabling the extrusion of Cl^-^ and maintaining KCC2 activity (Lee et al., 2007). Consistent with an earlier report (Liao and Lee, 2023), we observed a significant reduction in phosphorylation at the S940 site on KCC2 in the whole brain and forebrain lysates from *Cdkl5* KO mice. Additionally, we noted a significant increase in phosphorylation at the threonine 1007 and 906 sites on KCC2 (pKCC2-T1007), which inhibits its transport function (Moore et al., 2018). These findings are in line with literature indicating that the serine and threonine kinomes are altered in *Cdkl5* KO mice (Wang et al., 2012). This alteration may result from the inactivation of the mTOR signaling pathway (Wang et al., 2012), which is essential for the phosphorylation of KCC2 (Huang et al., 2012). Phosphorylation at these sites is necessary for the switch of GABA_A_-mediated excitation to inhibition during the second week of postnatal development. Our findings of altered phosphorylation suggest a potential delay in this critical developmental switch, which may contribute to hyperexcitability and infantile seizures in both human patients and animal models of CDD.

Neuronal and circuit hyperexcitability has been noted in EEG recordings from CDD patients (Saby et al., 2022), EEG and brain slice recordings from conditional *Cdkl5* knockout mice (Wang et al., 2020), as well as multielectrode recordings of neurons from CDD patient-derived organoids (Glass et al., 2024) To investigate this phenomenon, we compared baseline EEG power in *Cdkl5* KO mice to that of wild-type mice. Our analysis revealed a significant increase in baseline EEG power among the *Cdkl5* KO mice. This is in line with EEG studies conducted on human patients (Saby et al., 2022), where we observed elevated EEG power across the delta and theta frequency bands. These findings suggest that quantitative EEG measurements may serve as reliable biomarkers for diagnosing CDD and for the effective development of new treatments. Furthermore, we found that administering a KCC2 activator during postnatal development is sufficient to normalize EEG power levels in adult *Cdkl5* KO mice. Currently, there is a lack of validated biomarkers to assess brain function and clinical severity in individuals with CDD, which complicates the objective evaluation of emerging treatments. Therefore, we propose the use of quantitative EEG parameters as objective measures of brain function and disease severity in future clinical trials for CDD.

In rodent studies, SE induced by KA has been shown to be resistant to termination by diazepam, modeling patients experiencing drug-resistant seizures (Bertoglio et al.; Sharma et al., 2018; Drysdale et al., 2021; Colmers et al., 2024). Our research indicates that *Cdkl5* KO mice exhibit a shorter time to first seizure and develop diazepam-resistant SE more quickly than WT mice. Furthermore, the *Cdkl5* KO mice tend to spend more time engaged in epileptic activity, potentially due to their elevated basal EEG power. Notably, we discovered that repeated administration of OV350 during postnatal development was sufficient in delaying the onset of diazepam-resistant seizures and SE in adult *Cdkl5* KO mice.

Major autistic-like phenotypes, such as social and communication deficits, are hallmark characteristics of CDD (Okuda et al., 2018; Tang et al., 2019; Jhang et al., 2017; Jhang 2020). In line with existing literature, our findings indicate that *Cdkl5* KO mice performed significantly worse in social interaction tasks than their wild-type counterparts. We observed an alleviation of sociability deficits when we enhanced KCC2 activity in *Cdkl5* KO mice during postnatal development using OV350. However, activating KCC2 in adulthood did not lead to a reduction in these social deficits. This underscores the importance of early intervention during postnatal development, as this period is crucial for forming neural circuits and the accumulation of KCC2, which facilitates the transition from excitation to inhibition in immature neurons.

Mice that lack CDKL5 exhibit several key characteristics associated with the neurological disorder, including impairments in hippocampal-dependent memory and deficits in motor coordination. While both young adult and middle-aged *Cdkl5* KO mice demonstrate significant learning and memory deficits, they do not present motor impairments at an early age. Consistent with these observations, our studies revealed impairments in spatial learning and memory retention among *Cdkl5* KO mice. We found that enhancing KCC2 activity during the postnatal development of these *Cdkl5* KO mice led to long-term improvements in hippocampus-dependent spatial learning and memory retention. As a result, these mice performed better in learning the Barnes maze task and exhibited improved short-term memory. However, the enhancement of KCC2 activity during adulthood did not improve spatial memory as effectively as when treatment was given during infancy. This aligns with our previous research, which indicated that the constitutive activation of KCC2 through the development of a transgenic mouse line enhanced spatial learning and memory. Overall, our findings reaffirm the notion that early interventions may be the most effective strategy to reduce cognitive deficits in *Cdkl5* KO mice.

In summary, we observed that the loss of CDKL5 results in aberrant phosphorylation of KCC2. We also found that CDKL5 null mice have higher baseline EEG power, are more susceptible to kainate-induced, drug-resistant seizures, and show poor sociability and deficits in spatial learning and memory compared to WT mice. These deficits in adult mice are negated by potentiating KCC2 function during a period of neurodevelopment that is critical for circuit connectivity. This study demonstrates that enhancing KCC2 function during infancy is a potential therapy for CDD and other developmental and epileptic encephalopathies.

## Acknowledgments

This work was supported by Ovid Therapeutics, Inc., This study also received support from National Institutes of Health (NIH), National Institute of Neurological Disorders and Stroke Grants NS081986 (SM), NS101888 (SM), NS103865 (SM), NS111338 (SM), NS108378 (PD and SM), R21NS126914 (PD), R21NS111338 (PD), and R21NS111064 (PD), NIH National Institute of Mental Health Grant MH097446 (PD and SM).

---

**Supplementary Figure 1.**
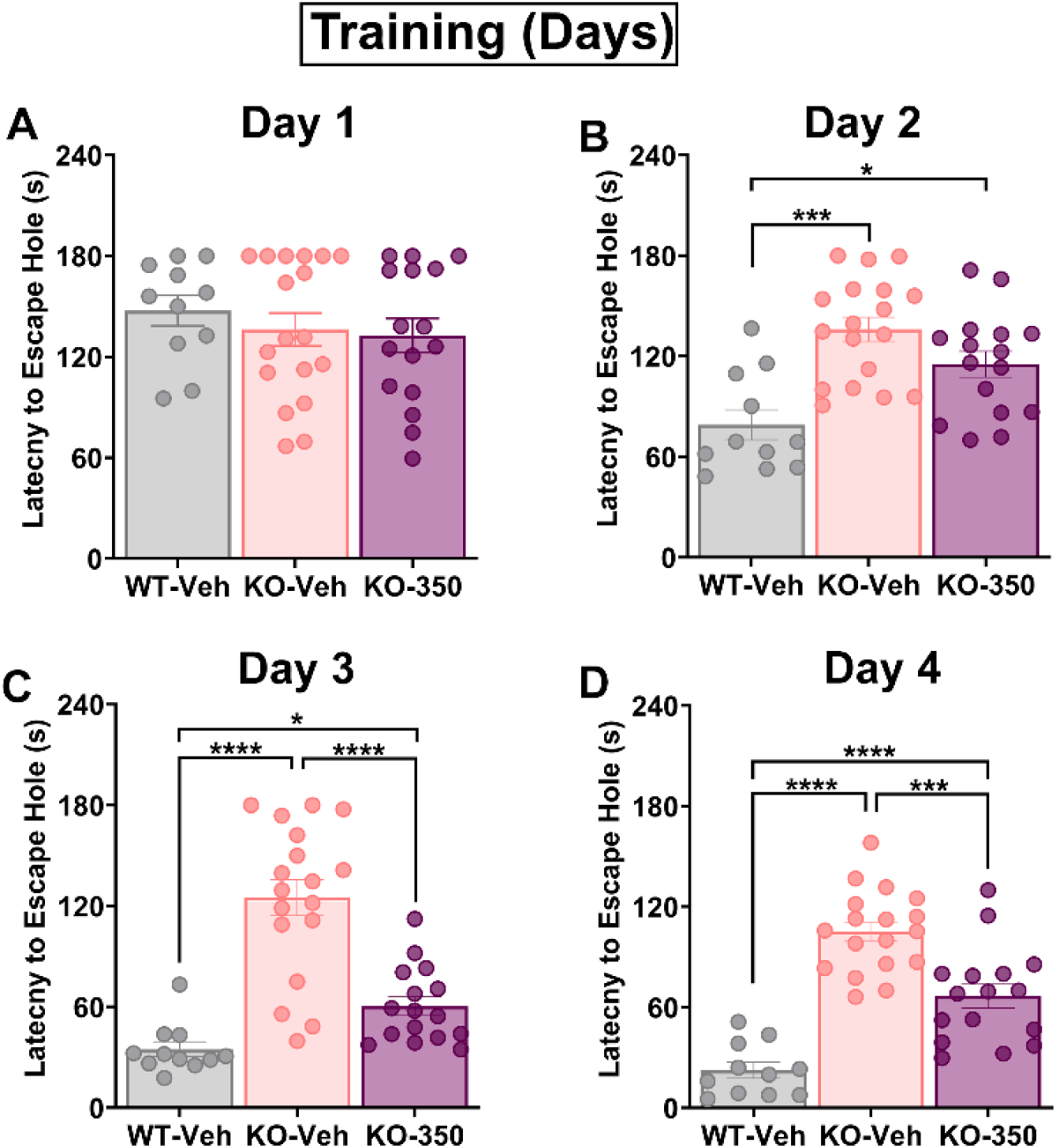
Potentiating KCC2 activity during development improves spatial learning in *Cdkl5* KO mice. Mice were exposed to the maze for four consecutive days (three training sessions each day), learning to associate the location of an escape hole using training room spatial cues. **A)** The latency to enter the escape hole was similar among the three treatment groups. **B)** The WT mice took less time to enter the escape hole than vehicle and OV350-treated (50 mg/kg *i.p*.) *Cdkl5*-KO mice on day 2. **C)** On day 3, OV350-treated *Cdkl5* KO mice showed significant improvement in spatial learning compared to vehicle-treated *Cdkl5* KO mice but didn’t bring it to the WT level. **D)** On day 4, OV350-treated *Cdkl5* KO mice showed further improvement in spatial learning compared to vehicle-treated *Cdkl5* KO mice but didn’t bring it to the WT level. The asterisk indicates the significant difference in latency to the escape hole. See table 2 for statistical analysis.

**Supplementary Figure 2.**
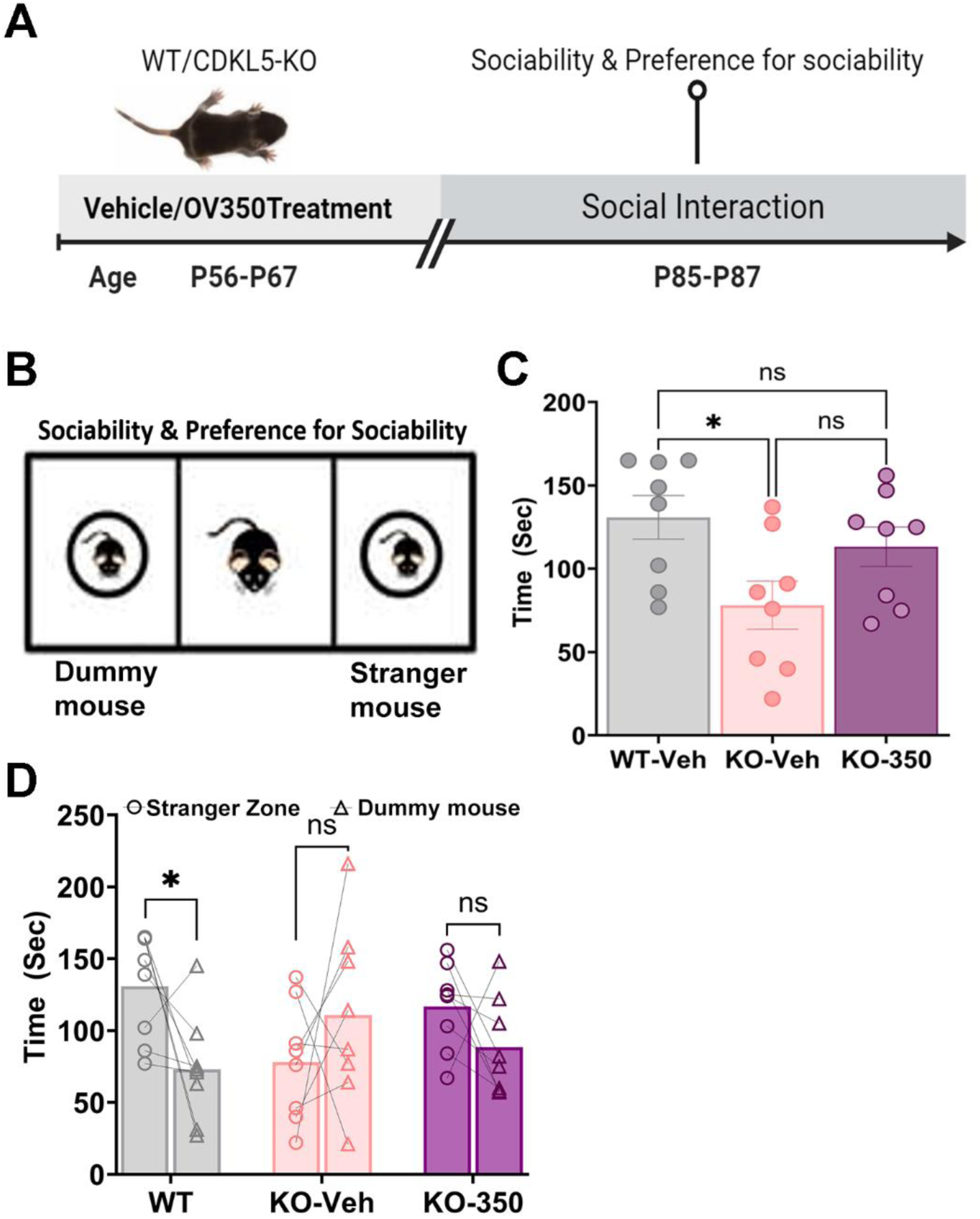
KCC2 activation during adulthood has no impact on social behavior. **A)** Experimental timeline. Seven to eight-week-old CDD mice were treated with either vehicle or OV350 (50 mg/kg *i.p*.). However, wild-type received vehicle injections only. **B)** Diagram of the sociability assay. Mice were allowed to explore either an unfamiliar mouse or a dummy mouse. **C)** Wild-type mice spent more time interacting with the mouse than vehicle-treated *Cdkl5* KO mice, and no differences were observed between the other groups. The asterisk indicates the significance of the time spent with the stranger mouse. **D)** WT mice preferred interacting with the mouse vs. the dummy mouse. No effects of OV350 were observed on sociability in *Cdkl5* KO mice. The asterisk indicates the significance of the time spent with the dummy and the stranger mouse. See table 2 for statistical analysis.

**Supplementary Figure 3.**
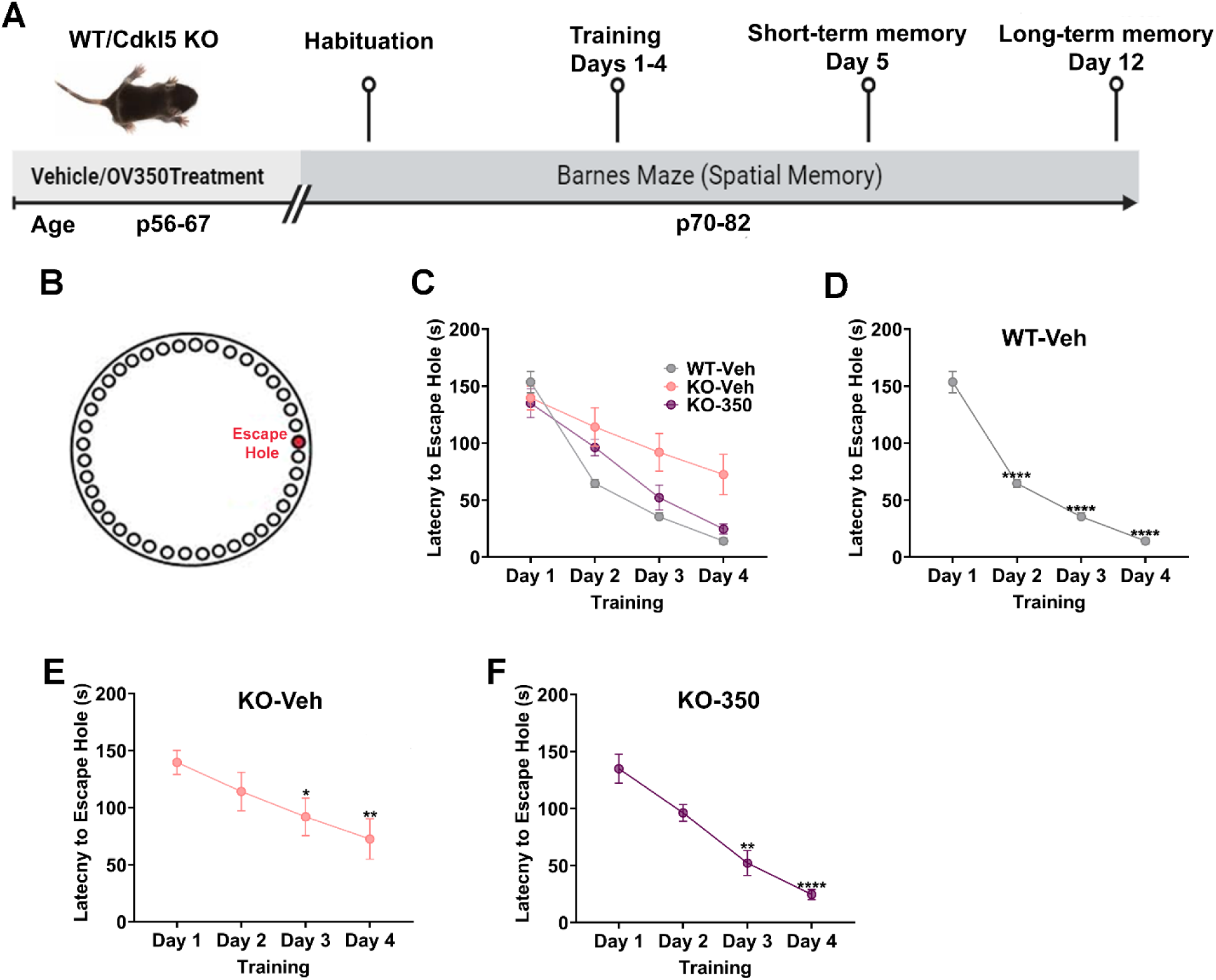
KCC2 activation during adulthood improves the rate of spatial learning in *Cdkl5* KO mice. **A)** Experimental timeline. Seven to eight-week-old *Cdkl5* KO mice were treated with either vehicle or OV350 (50 mg/kg *i.p*.). However, WT received vehicle injections only. **B)** Cartoon illustration of the Barnes maze. **C)** Latency to enter the goal hole was measured on days 1–4; wild-type mice learned better than the vehicle-treated *Cdkl5* KO mice from days 2-4. However, OV350-treated *Cdkl5* KO mice performed considerably better than vehicle-treated *Cdkl5* KO mice from days 3-4. **D)** The rate of learning (day when there is a significant reduction in latency to goal compared to day 1) was also comparable in wild-type mice. The asterisk indicates the significant difference in latency to the escape hole with reference to day 1. **E)** The learning rate was comparable in vehicle-treated *Cdkl5* KO mice from day 3. The asterisk indicates the significant difference in latency to the escape hole with reference to day 1. **F)** The learning rate was comparable in OV350-treated *Cdkl5* KO mice from day 3. The asterisk indicates the significant difference in latency to the escape hole with reference to day 1. See table 2 for statistical analysis.

**Supplementary Figure 4.**
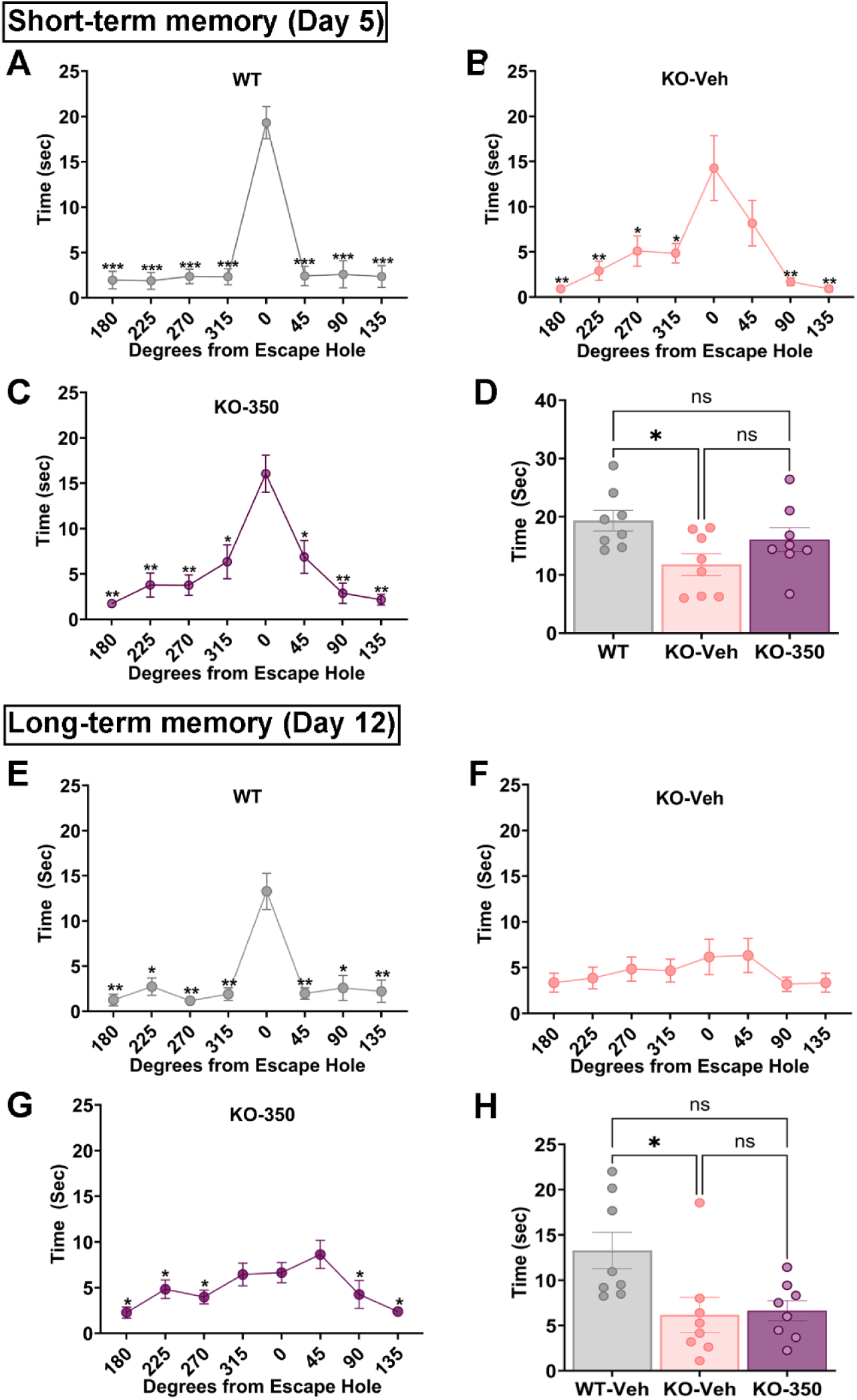
KCC2 activation during adulthood in *Cdkl5* KO mice does not affect spatial memory retention. Spatial memory was assessed using a Barnes maze assay. After 4 days of training, time spent at each hole was measured on day 5 and day 12 upon removal of the escape tunnel, and data was binned into 45° groups. **A)** Wild-type mice preferred the goal area on day 5. Vehicle-treated *Cdkl5* KO mice didn’t perform as well as the WT mice. OV350-treated (50 mg/kg *i.p*.) *Cdkl5* KO mice performed considerably well. Wild-type mice spent more time at the goal zone than the vehicle but not compared to the OV350-treated *Cdkl5* KO mice. B) WT mice show enhanced specificity of the spatial memory on day 12 compared to the vehicle and OV350-treated *Cdkl5* KO mice. The WT mice spent more time at the goal region than the other two groups of mice. * Indicates significant difference to time spent in the 0° region. See table 2 for statistical analysis.

## Notes

### Competing Interest Statement

S.J.M. serves as a consultant for AstraZeneca, Ovid Therapeutics, and Sage Therapeutics, relationships that are regulated by Tufts University.
S.J.M. holds equity in Sage Therapeutics. S.F.J.N., and S.J.M. hold equity in Ovid Therapeutics. T.N, and Z.Z. are employees of Ovid Therapeutics and hold equity.

## References

Amendola E, Zhan Y, Mattucci C, Castroflorio E, Calcagno E, Fuchs C, Lonetti G, Silingardi D, Vyssotski AL, Farley D, Ciani E, Pizzorusso T, Giustetto M, Gross CT (2014) Mapping Pathological Phenotypes in a Mouse Model of CDKL5 Disorder. PLOS ONE 9:e91613.

Antrobus R, Borner GHH (2011) Improved Elution Conditions for Native Co-Immunoprecipitation. PLOS ONE 6:e18218.

Arshad MN, Naegele JR (2020) Induction of Temporal Lobe Epilepsy in Mice with Pilocarpine. Bio Protoc 10:e3533.

Bengzon J, Kokaia Z, Elmér E, Nanobashvili A, Kokaia M, Lindvall O (1997) Apoptosis and proliferation of dentate gyrus neurons after single and intermittent limbic seizures. Proceedings of the National Academy of Sciences 94:10432–10437.

Bertoglio D, Amhaoul H, Van Eetveldt A, Houbrechts R, Van De Vijver S, Ali I, Dedeurwaerdere S (2017) Kainic Acid-Induced Post-Status Epilepticus Models of Temporal Lobe Epilepsy with Diverging Seizure Phenotype and Neuropathology. Front Neurol 8:588.

Choi C, Smalley JL, Lemons AHS, Ren Q, Bope CE, Dengler JS, Davies PA, Moss SJ (2022) Analyzing the mechanisms that facilitate the subtype-specific assembly of γ-aminobutyric acid type A receptors. Front Mol Neurosci 15:1017404.

Colmers PLW, Arshad MN, Mukherjee J, Lin S, Ng SFJ, Sarmiere P, Davies PA, Moss SJ (2024) Sustained Inhibition of GABA-AT by OV329 Enhances Neuronal Inhibition and Prevents Development of Benzodiazepine Refractory Seizures. eNeuro 11.

D’Mello SR (2023) Rett and Rett-related disorders: Common mechanisms for shared symptoms? Exp Biol Med (Maywood) 248:2095–2108.

Deeb TZ, Maguire J, Moss SJ (2012) Possible alterations in GABAA receptor signaling that underlie benzodiazepine-resistant seizures. Epilepsia 53:79–88.

Deshpande LS, Blair RE, Nagarkatti N, Sombati S, Martin BR, DeLorenzo RJ (2007) Development of pharmacoresistance to benzodiazepines but not cannabinoids in the hippocampal neuronal culture model of status epilepticus. Exp Neurol 204:705–713.

Duarte ST, Armstrong J, Roche A, Ortez C, Pérez A, O’Callaghan Mdel M, Pereira A, Sanmartí F, Ormazábal A, Artuch R, Pineda M, García-Cazorla A (2013) Abnormal expression of cerebrospinal fluid cation chloride cotransporters in patients with Rett syndrome. PLoS One 8:e68851.

Fuchs C, Gennaccaro L, Trazzi S, Bastianini S, Bettini S, Lo Martire V, Ren E, Medici G, Zoccoli G, Rimondini R, Ciani E (2018) Heterozygous CDKL5 Knockout Female Mice Are a Valuable Animal Model for CDKL5 Disorder. Neural Plast 2018:9726950.

Furukawa T, Fukuda A (2023) Maternal taurine as a modulator of Cl(-) homeostasis as well as of glycine/GABA(A) receptors for neocortical development. Front Cell Neurosci 17:1221441.

Glass MR, Whye D, Anderson NC, Wood D, Makhortova NR, Polanco T, Kim KH, Donovan KE, Vaccaro L, Jain A, Cacchiarelli D, Sun L, Olson H, Buttermore ED, Sahin M (2024) Excitatory Cortical Neurons from CDKL5 Deficiency Disorder Patient-Derived Organoids Show Early Hyperexcitability Not Identified in Neurogenin2 Induced Neurons. bioRxiv:2024.2011.2011.622878.

Goodkin HP, Joshi S, Mtchedlishvili Z, Brar J, Kapur J (2008) Subunit-specific trafficking of GABA(A) receptors during status epilepticus. J Neurosci 28:2527–2538.

He Q, Nomura T, Xu J, Contractor A (2014) The developmental switch in GABA polarity is delayed in fragile X mice. J Neurosci 34:446–450.

Hu Z, Yu X, Chen P, Jin K, Zhou J, Wang G, Yu J, Wu T, Wang Y, Lin F, Zhang T, Wang Y, Zhao X (2023) BDNF-TrkB signaling pathway-mediated microglial activation induces neuronal KCC2 downregulation contributing to dynamic allodynia following spared nerve injury. Mol Pain 19:17448069231185439.

Huang X, McMahon J, Yang J, Shin D, Huang Y (2012) Rapamycin down-regulates KCC2 expression and increases seizure susceptibility to convulsants in immature rats. Neuroscience 219:33–47.

Jarvis R et al. (2023) Direct activation of KCC2 arrests benzodiazepine refractory status epilepticus and limits the subsequent neuronal injury in mice. Cell Rep Med 4:100957.

Jhang C-L, Huang T-N, Hsueh Y-P, Liao W (2017) Mice lacking cyclin-dependent kinase-like 5 manifest autistic and ADHD-like behaviors. Human Molecular Genetics 26:3922–3934.

Kelley MR, Deeb TZ, Brandon NJ, Dunlop J, Davies PA, Moss SJ (2016) Compromising KCC2 transporter activity enhances the development of continuous seizure activity. Neuropharmacology 108:103–110.

Knight EMP et al. (2022) Safety and efficacy of ganaxolone in patients with CDKL5 deficiency disorder: results from the double-blind phase of a randomised, placebo-controlled, phase 3 trial. The Lancet Neurology 21:417–427.

Laxer KD, Trinka E, Hirsch LJ, Cendes F, Langfitt J, Delanty N, Resnick T, Benbadis SR (2014) The consequences of refractory epilepsy and its treatment. Epilepsy Behav 37:59–70.

Lee HH, Walker JA, Williams JR, Goodier RJ, Payne JA, Moss SJ (2007) Direct protein kinase C-dependent phosphorylation regulates the cell surface stability and activity of the potassium chloride cotransporter KCC2. J Biol Chem 282:29777–29784.

Leonard H, Downs J, Benke TA, Swanson L, Olson H, Demarest S (2022) CDKL5 deficiency disorder: clinical features, diagnosis, and management. Lancet Neurol 21:563–576.

Liao W, Lee K-Z, Su S-H, Luo Y (2020) Deficiency of cyclin-dependent kinase-like 5 causes spontaneous epileptic seizures in neonatal mice. bioRxiv:2020.2003.2009.983981.

McMoneagle E, Zhou J, Zhang S, Huang W, Josiah SS, Ding K, Wang Y, Zhang J (2024) Neuronal K+- Cl- cotransporter KCC2 as a promising drug target for epilepsy treatment. Acta Pharmacologica Sinica 45:1–22.

Moore YE, Kelley MR, Brandon NJ, Deeb TZ, Moss SJ (2017) Seizing Control of KCC2: A New Therapeutic Target for Epilepsy. Trends Neurosci 40:555–571.

Moore YE, Deeb TZ, Chadchankar H, Brandon NJ, Moss SJ (2018) Potentiating KCC2 activity is sufficient to limit the onset and severity of seizures. Proc Natl Acad Sci U S A 115:10166–10171.

Moore YE, Conway LC, Wobst HJ, Brandon NJ, Deeb TZ, Moss SJ (2019) Developmental Regulation of KCC2 Phosphorylation Has Long-Term Impacts on Cognitive Function. Front Mol Neurosci 12:173.

Okuda K, Takao K, Watanabe A, Miyakawa T, Mizuguchi M, Tanaka T (2018) Comprehensive behavioral analysis of the Cdkl5 knockout mice revealed significant enhancement in anxiety- and fear-related behaviors and impairment in both acquisition and long-term retention of spatial reference memory. PLOS ONE 13:e0196587.

Pizzo R, Gurgone A, Castroflorio E, Amendola E, Gross C, Sassoè-Pognetto M, Giustetto M (2016) Lack of Cdkl5 Disrupts the Organization of Excitatory and Inhibitory Synapses and Parvalbumin Interneurons in the Primary Visual Cortex. Front Cell Neurosci 10:261.

Rattka M, Brandt C, Löscher W (2013) The intrahippocampal kainate model of temporal lobe epilepsy revisited: epileptogenesis, behavioral and cognitive alterations, pharmacological response, and hippoccampal damage in epileptic rats. Epilepsy Res 103:135–152.

Reh R, Williams LJ, Todd RM, Ward LM (2021) Warped rhythms: Epileptic activity during critical periods disrupts the development of neural networks for human communication. Behavioural Brain Research 399:113016.

Rivera C, Li H, Thomas-Crusells J, Lahtinen H, Viitanen T, Nanobashvili A, Kokaia Z, Airaksinen MS, Voipio J, Kaila K, Saarma M (2002) BDNF-induced TrkB activation down-regulates the K+-Cl- cotransporter KCC2 and impairs neuronal Cl- extrusion. J Cell Biol 159:747–752.

Saby JN, Mulcahey PJ, Zavez AE, Peters SU, Standridge SM, Swanson LC, Lieberman DN, Olson HE, Key AP, Percy AK, Neul JL, Nelson CA, Roberts TPL, Benke TA, Marsh ED (2022) Electrophysiological biomarkers of brain function in CDKL5 deficiency disorder. Brain Commun 4:fcac197.

Sharma AK, Jordan WH, Reams RY, Hall DG, Snyder PW (2008) Temporal profile of clinical signs and histopathologic changes in an F-344 rat model of kainic acid-induced mesial temporal lobe epilepsy. Toxicol Pathol 36:932–943.

Silayeva L, Deeb TZ, Hines RM, Kelley MR, Munoz MB, Lee HH, Brandon NJ, Dunlop J, Maguire J, Davies PA, Moss SJ (2015) KCC2 activity is critical in limiting the onset and severity of status epilepticus. Proc Natl Acad Sci U S A 112:3523–3528.

Sivakumaran S, Cardarelli RA, Maguire J, Kelley MR, Silayeva L, Morrow DH, Mukherjee J, Moore YE, Mather RJ, Duggan ME, Brandon NJ, Dunlop J, Zicha S, Moss SJ, Deeb TZ (2015) Selective inhibition of KCC2 leads to hyperexcitability and epileptiform discharges in hippocampal slices and in vivo. J Neurosci 35:8291–8296.

Smalley JL, Kontou G, Choi C, Ren Q, Albrecht D, Abiraman K, Santos MAR, Bope CE, Deeb TZ, Davies PA, Brandon NJ, Moss SJ (2020) Isolation and Characterization of Multi-Protein Complexes Enriched in the K-Cl Co-transporter 2 From Brain Plasma Membranes. Frontiers in Molecular Neuroscience 13.

Smalley JL, Cho N, Ng SFJ, Choi C, Lemons AHS, Chaudry S, Bope CE, Dengler JS, Zhang C, Rasband MN, Davies PA, Moss SJ (2023) Spectrin-beta 2 facilitates the selective accumulation of GABAA receptors at somatodendritic synapses. Communications Biology 6:11.

Symonds JD, McTague A (2020) Epilepsy and developmental disorders: Next generation sequencing in the clinic. Eur J Paediatr Neurol 24:15–23.

Tang X, Drotar J, Li K, Clairmont CD, Brumm AS, Sullins AJ, Wu H, Liu XS, Wang J, Gray NS, Sur M, Jaenisch R (2019) Pharmacological enhancement of KCC2 gene expression exerts therapeutic effects on human Rett syndrome neurons and Mecp2 mutant mice. Sci Transl Med 11.

Tipton AE, Russek SJ (2022) Regulation of Inhibitory Signaling at the Receptor and Cellular Level; Advances in Our Understanding of GABAergic Neurotransmission and the Mechanisms by Which It Is Disrupted in Epilepsy. Front Synaptic Neurosci 14:914374.

Virtanen MA, Uvarov P, Mavrovic M, Poncer JC, Kaila K (2021) The Multifaceted Roles of KCC2 in Cortical Development. Trends in Neurosciences 44:378–392.

Wang HT, Zhu ZA, Li YY, Lou SS, Yang G, Feng X, Xu W, Huang ZL, Cheng X, Xiong ZQ (2021) CDKL5 deficiency in forebrain glutamatergic neurons results in recurrent spontaneous seizures. Epilepsia 62:517–528.

Wang IT, Allen M, Goffin D, Zhu X, Fairless AH, Brodkin ES, Siegel SJ, Marsh ED, Blendy JA, Zhou Z (2012) Loss of CDKL5 disrupts kinome profile and event-related potentials leading to autistic-like phenotypes in mice. Proc Natl Acad Sci U S A 109:21516–21521.

Wong K, Junaid M, Alexander S, Olson HE, Pestana-Knight EM, Rajaraman RR, Downs J, Leonard H (2024) Caregiver Perspective of Benefits and Side Effects of Anti-Seizure Medications in CDKL5 Deficiency Disorder from an International Database. CNS Drugs 38:719–732.

